# Reward expectation selectively boosts the firing of accumbens D1+ neurons during motivated approach

**DOI:** 10.1101/2023.09.02.556060

**Authors:** Thomas W. Faust, Ali Mohebi, Joshua D. Berke

## Abstract

The nucleus accumbens (NAc) helps govern motivation to pursue rewards. Two distinct sets of NAc projection neurons-expressing dopamine D1 versus D2 receptors-are thought to promote and suppress motivated behaviors respectively. However, support for this conceptual framework is limited: in particular the spiking patterns of these distinct cell types during motivated behavior have been largely unknown. We monitored identified D1+ and D2+ neurons in the NAc Core, as unrestrained rats performed an operant task in which motivation to initiate work tracks recent reward rate. D1+ neurons preferentially increased firing as rats initiated trials, and fired more when reward expectation was higher. By contrast, D2+ cells preferentially increased firing later in the trial especially in response to reward delivery - a finding not anticipated from current theoretical models. Our results provide new evidence for the specific contribution of NAc D1+ cells to self-initiated approach behavior, and will spur updated models of how we learn from rewards.

## Introduction

Many rewards can only be obtained with time and effort, and even then success is uncertain. Deciding whether rewards are worth working for is a key function of brain circuits involving the nucleus accumbens (NAc), a ventral-medial portion of the striatum. Alterations in NAc information processing are implicated in a range of human behavioral disorders, including drug addiction (1) and depression (2), and the NAc is a major target for emerging neuromodulation therapies for these conditions (3).

Yet how exactly NAc processes information and influences downstream structures is poorly understood. As in the rest of striatum, the great majority (95%) of NAc cells are GABAergic spiny projection neurons (SPNs). These can be divided into two similarly sized subpopulations based on projection targets and gene expression (4) - in particular, they bear distinct dopamine (DA) receptors: D1r versus D2r (5). These SPN subpopulations (“D1+”, “D2+”) are thereby differently affected by NAc DA, a powerful modulator of both learning and motivation: animals’ willingness to work for rewards (6–8). DA has complex effects on striatal neurons (9) but - crudely - is thought to enhance excitability of D1+SPNs (10) and reduce excitability of D2+ SPNs (11). Partly because of these opposing acute DA effects, D1+ SPNs are often considered to promote behavior (“Go” pathway) while D2+ SPNs suppress behavior (“NoGo” pathway) (reviewed in (12)). At least in dorsal striatum, this scheme is supported by evidence from optogenetic manipulations of D1+ and D2+ cells (13).

Fluctuations in DA also control plasticity of corticostriatal synapses, in a distinct manner for D1+ and D2+ SPNs (14). Brief (“phasic”) increases in DA can encode reward prediction error (RPE; (15–17)): a message that reward predictions need to be updated (18). These DA pulses increase DA binding to D1r, activating cAMP/PKA-dependent signals to achieve persistent changes in synaptic strength and behavior (19, 20). Through this sensitivity to better-than-expected events, D1+ SPNs may subsequently enhance predictions of reward and thereby encourage behavior. D2+ SPNs are thought to have a complementary role. D2r are negatively coupled to cAMP/PKA pathways, so transient dips in DA may disinhibit plasticity mechanisms in D2+ SPNs (21). Through sensitivity to worse-than-expected events, D2+ SPNs may contribute to subsequent predictions that rewards will not be received, and discourage behavior (22).

However, in NAc this elegant theoretical framework has received only mixed support from manipulation and recording studies. For example, brief optogenetic activation of either NAc D1+ or D2+ cells was reported to increase motivation in a progressive-ratio test (23), and to produce a conditioned place preference (24). Investigations of individual NAc neurons in behaving animals tend to find a wide variety of firing patterns (25–28), and recent studies using calcium imaging suggest that this variability persists even after distinguishing D1+ and D2+ neurons (e.g. (29, 30)). Spiking patterns of identified D1+ and D2+ neurons have been studied to a limited extent in other striatal subregions (31, 32). Yet there is virtually no information available about the spiking of identified NAc neurons, and their relationships to motivation.

To overcome this gap and test key theories of NAc function, we recorded the spiking of substantial numbers of optogenetically identified, individual NAc D1+ and D2+ SPNs in freely behaving rats. We took advantage of an operant behavioral task in which rats’ willingness to work depends strongly on whether or not recent trials have been rewarded, and in which we have extensively characterized NAc DA signals (17, 33).

We examined activity throughout task performance, but our hypotheses focused primarily on two time epochs. The first is when rats decide to initiate trials by approaching a nose poke port. This motivated approach behavior is accompanied by a brief ramping increase in NAc DA, and in similar contexts is dependent on NAc D1r (34). We hypothesized that NAc D1+ neurons would be particularly involved during this approach, and their activity would positively scale with value (i.e. higher when recent trials have been rewarded). Our results confirm this hypothesis. The second key time epoch is shortly after reward delivery (or omission). We previously reported that the sound of reward delivery evokes a marked increase in NAc DA, that scales with RPE (i.e. it is stronger when recent trials have not been rewarded), and that omission produces a DA dip (17, 33). We hypothesized that D1+ and D2+ SPNs would preferentially increase firing to rewards and omissions respectively. Contrary to this hypothesis, we found greater activation of NAc D2+ SPNs at reward delivery.

## Results

### Optogenetic identification of D1+ and D2+ SPNs during operant task performance

To distinguish SPN sub-populations we made use of two transgenic knockin rat lines, D1-Cre and A2a-Cre. As we previously described in detail (35), these produce faithful expression of Cre recombinase in D1+ and D2+ neurons respectively. We used A2a-Cre instead of D2-Cre rats because D2r is also expressed by e.g. cholinergic interneurons (36), while expression of A2a receptors is selective to D2+ SPNs (37). In each rat we bilaterally infused virus into NAc for Cre-dependent expression of the red-light-sensitive opsin ChrimsonR ((38); see Methods). We implanted a chronic recording device (Fig. 1A) consisting of two optic fibers (one per hemisphere) each surrounded by 16 tetrodes (4-wire bundles; 128 wires total/rat). We targeted the NAc Core, which has been particularly implicated in value-guided decisions to work (39). Histological analyses confirmed that the great majority of our recording locations were within the Core (Fig. 1B, C). At the end of each recording session we applied a series of brief (10ms) pulses of red light through the optic fibers. Neurons were considered “tagged” as light-responsive - and therefore either D1+, or A2a+ and D2+, depending on rat line - if they showed a significant increase in firing within 10ms of light onset (Fig. 1D-F; see Methods for full criteria).

**Fig. 1.**
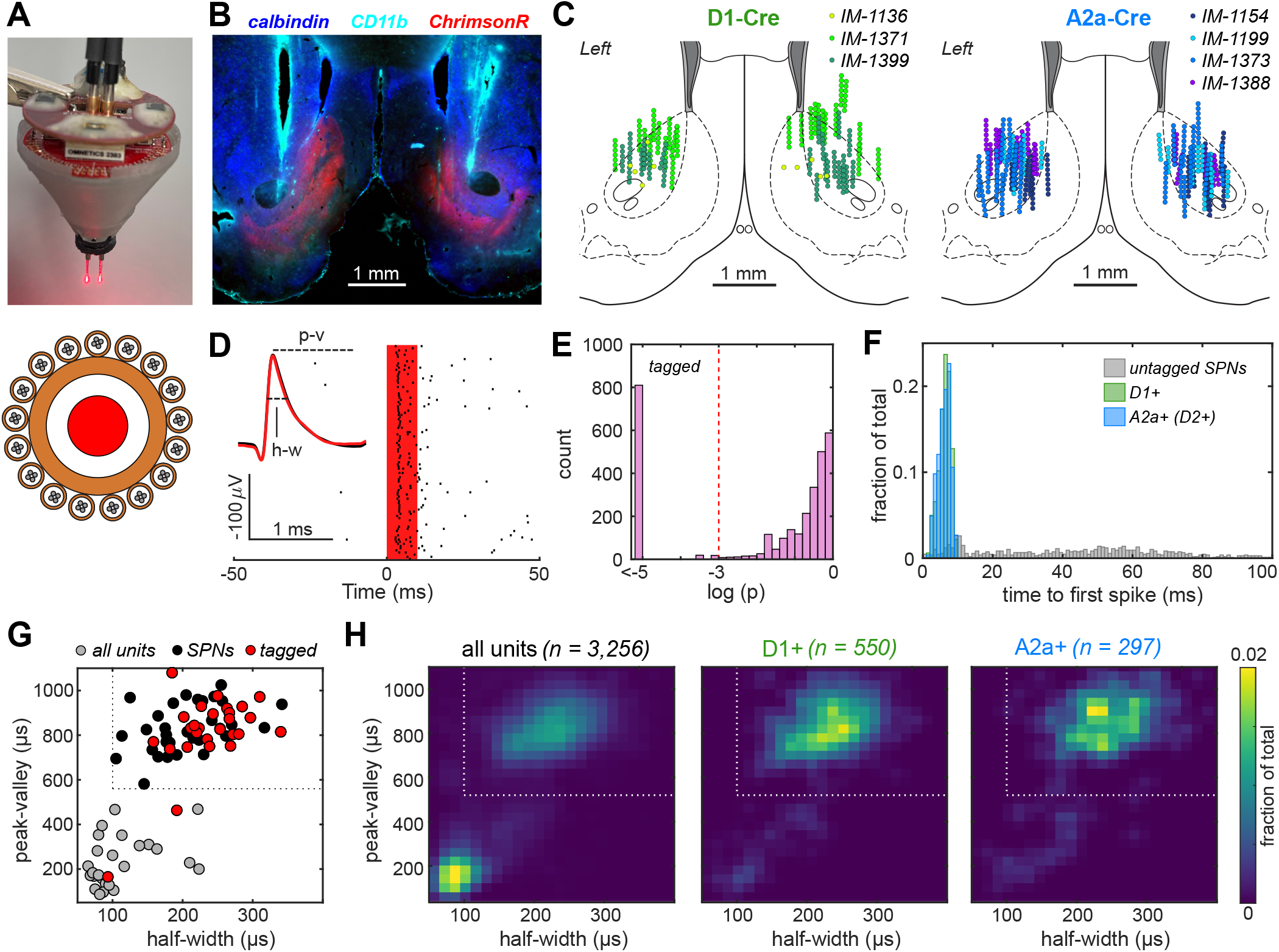
Opto-tagging of D1+ and D2+ SPNs. **A**, Top, drive assembly; bottom, schematic of implanted bundle (one per side) of tetrodes surrounding an optic fiber. **B**, Histology example (coronal section). Staining for CD11b (cyan) marks damage caused by the electrodes; calbindin distinguishes the NAc Core (blue) from unlabeled Shell, and ChrimsontdTomato (red) shows area of transfected neurons (D1+ in this case). **C**, Recording locations reconstructed from tetrode marks, with individual D1-Cre and A2a-Cre rats shown by different colors. **D**, Example spiking of one tagged D1+ SPN to red laser pulses (10 ms duration, 100 trials); inset shows average waveforms for spontaneous spikes (black) and laser-evoked spikes (red). Spike half-width (h-w) and peak-valley duration (p-v) for this cell are marked. **E**, Distribution of tagging significance p levels for all recorded units (see Methods), indicating p = 0.001 threshold. **F**, Average time to first spike following the onset of 10 ms laser pulses for tagged D1+ and D2+ units and untagged SPNs. **G**, Distribution of spike waveform features of all recorded units in one example session, with presumed SPNs in black, other cells (presumed interneurons) in grey, and light-responsive neurons in red. Dotted lines indicate boundaries used to separate SPNs (half-width > 100 *μ*s, peak-valley duration > 560 *μ*s). **H**, Density of units in waveform feature-space for all recorded units (left), D1-Cre tagged units (center), and A2a-Cre tagged units (right).

We isolated 3,256 distinct single neurons from 44 recording sessions in 7 rats (3 D1-Cre, 4 A2a-Cre). Examination of waveform shape produced two major clusters (Fig. 1G, H), consistent with prior studies performing extracellular recordings in behaving rodents (40–42). The smaller cluster with brief spike waveforms has higher firing rates and has been presumed to be interneurons, while the larger cluster with longer spike waveforms (60.2%; n = 1,961) has been presumed to represent spiny projection neurons (SPNs). Consistent with this, we found that the great majority of optotagged cells fell into this SPN cluster (Fig. 1H), for both D1+ (91.3%; n = 502) and A2a+ (89.2%; n = 265) neurons. All of our subsequent analyses focused on tagged cells within this SPN cluster, which for simplicity we refer to as D1+ and D2+ cells below. These subpopulations had similarly low average firing rates (session-wide medians: D1+ 0.27 Hz, D2+ 0.28 Hz; no difference, p = 0.401, permutation test).

During each recording session rats performed an operant ‘bandit’ task (Hamid et al., 2019; Mohebi et al., 2019). In brief, illumination of a central hole (‘Light-On’) indicates that the rat can initiate a trial by nose-poking that hole (‘Center-In’). If the rat maintains this nose poke for the full duration of a variable hold period (500-1500 ms), it hears a white noise burst (‘Go Cue’), prompting movement out of the central hole (‘Center-Out’) and into a hole immediately to the left or right (‘Side-In’). At this point the rat may (or may not) hear the click sound of a food hopper delivering a sucrose reward pellet at another location behind them, which the rat can then collect (‘Food-Port-In’). Reward probabilities for leftward and rightward choices (10, 50, 90%) were independently and pseudo-randomly adjusted after each block of 35-45 trials (Fig. 2A). All rats received extensive pretraining with these task parameters (>3 months) prior to implantation.

Through trial-and-error, rats adjusted both their left/right choices, and their motivation to perform the task (Fig. 2A-C). In particular, their mean “latency” (time between Light-On and Center-In) was shorter when they had higher expectation of reward, due to a higher rate of recent rewards. To estimate this reward expectation we used a simple trial-based reinforcement learning model: “trial value” (V) increased with each reward and decreased with each omission (between 0 and 1, recency-weighted; see Methods).

**Fig. 2.**
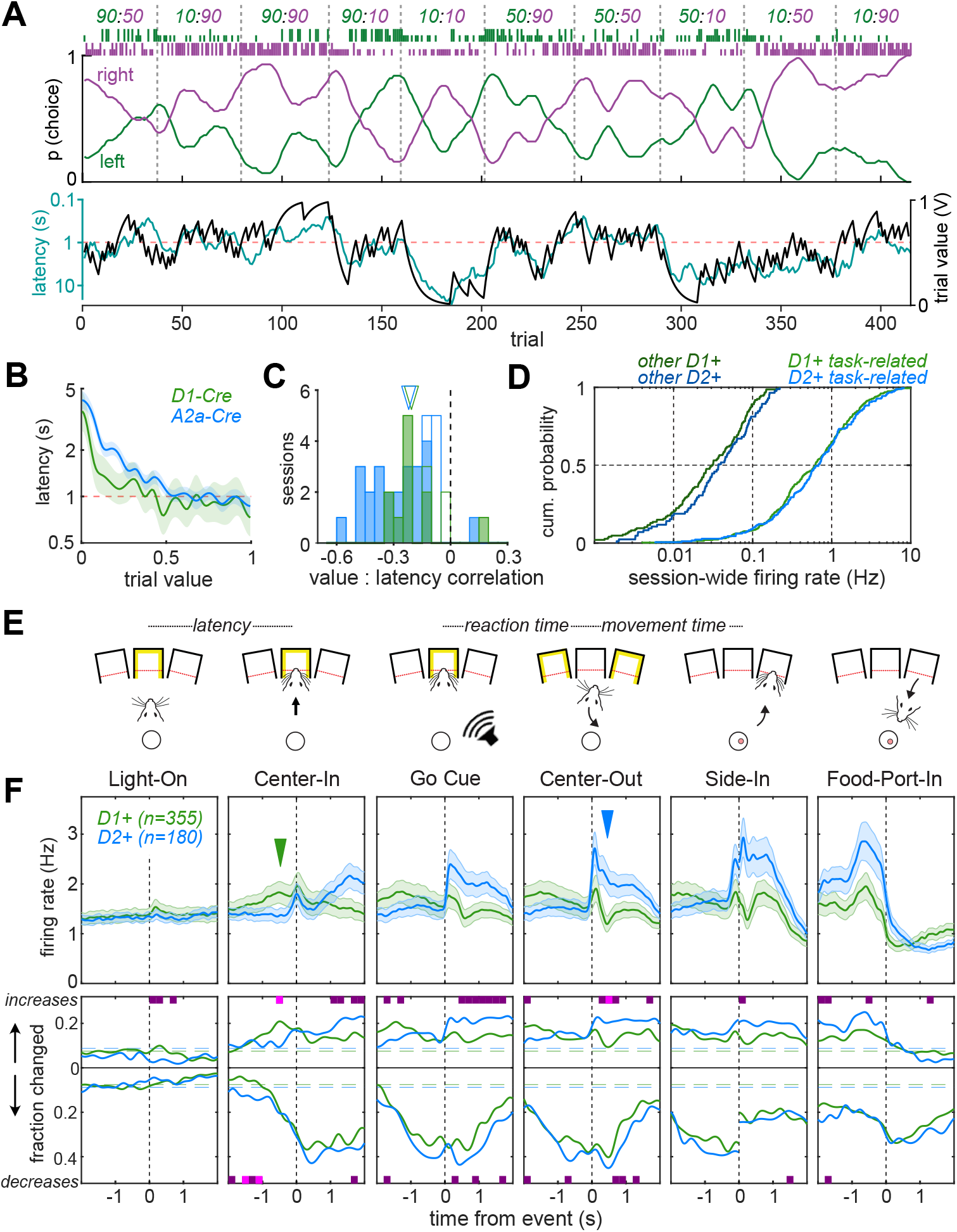
Distinct responses of D1+ and D2+ SPNs in the operant task. **A**, Example session. From top, reward probabilities for left (green) and right (purple) choices; tall ticks indicate rewarded trials, short ticks unrewarded trials; running averages of choices (10-trial smoothing). Bottom plots, estimated trial value (black), and running average of latency (cyan, 10-trial smoothing). **B**, Average latencies by trial value for D1-Cre and A2a-Cre sessions (mean +/-SEM, D1-Cre n = 14, A2a-Cre n = 29).**C**, Correlation coefficients between trial value and log(latency) for D1-Cre (green) and A2a-Cre (blue) sessions, with significant correlations in filled bars (p < 0.05). **D**, Cumulative distributions of session-wide firing rates for D1+ and D2+ SPNs, subdivided into task-related (D1+ n=355, D2+ n=180) and others (D1+ n=147, D2+ n=85). **E**, Behavioral event sequence. **F**, Event-aligned activity of task-related D1+ and D2+ subpopulations. Top, average firing rates (+/-SEM); bottom, fractions of each subpopulation with increased or decreased firing. Colored bars indicate time bins when these fractions were significantly different between D1+ and D2+ (p < 0.05, purple: uncorrected; magenta: Bonferroni-corrected). Blue, green arrowheads point to moments of most significant divergence between average activity of each population (D1+ > D2+ and D2+ > D1+, respectively). Around each event we include a time range of-2:+2s, the windows used to define “task-related” cells, but note that these windows overlap with each other to varying degrees on each trial. Data are from trials with latencies > 1s (i.e. “non-engaged”; see Fig. 3), and after Side-In only rewarded trials are included.

As expected from prior studies, many SPNs maintained low firing rates during task performance. We defined “task-related” cells as those that showed a significant increase or decrease relative to session-wide firing rate, near any of six behavioral events (Fig. 2E; one-tailed permutation tests using 200 ms bins; p<0.05, corrected for multiple comparisons). The majority of SPNs in each tagged subpopulation were task-related (D1+: 355/502, 70.7%; D2+: 180/265, 67.9%).These cells had higher mean firing rates compared to the remaining SPNs (D1+ and D2+ each p < 0.0001, permutation tests), although for virtually all D1+ and D2+ cells mean firing rates were below 10Hz (Fig. 2D). For reference we show the patterns of firing rate increases and decreases for each tagged task-related neuron in Supplementary Fig. 1).

### Distinct gross activity patterns of D1+ and D2+ SPNs around behavioral events

We next considered population activity around each behavioral event, assessed either as mean firing rate for each subpopulation (Fig. 2F, top), or the fractions of individual cells in each subpopulation showing significant increases or decreases in activity (Fig. 2F, bottom; fraction of cells undergoing significant increases or decreases in firing rate compared to permutated distribution). For both D1+ and D2+ SPNs, subsets of cells became more active during performance of each trial, and larger subsets (>40% of each subpopulation) became less active (Fig. 2F, bottom; Supplementary Fig. 1; see also (43)). Nonetheless we observed two obvious differences between D1+ and D2+ SPNs. Shortly before Center-In, there was a selective increase in D1+ firing (green arrowhead). Subsequently, after the onset of the Go cue there was a preferential increase in D2+ activity (blue arrowhead), and on rewarded trials this persisted until the trial was completed at Food-Port-In.

### Increased D1+ firing during motivated approach, in proportion to reward expectation

We examined in more detail the early, selective increase in D1+ activity. In this operant task we have previously observed (33) that in some trials, at the time of Light-On rats are already waiting near the center port (“engaged”; Fig. 3A), and can poke immediately resulting in very short latencies. In other trials rats are elsewhere in the chamber (“non-engaged”), and it takes time for them to approach the illuminated port to initiate the trial. Video analysis showed that these two trial types result in a bimodal distribution of shorter and longer latencies respectively ((33); Fig. 3B). The two modes separate at 1s, which is the approximate duration of the approach behavior (33). We tested how this engaged/non-engaged difference affects the subpopulations of D1+ and D2+ SPNs which increase firing during this part of the trial (defined as 4s before Center-In to 1s after). A ramping increase in D1+ SPNs was preferentially seen for non-engaged trials, as rats approached the Center port (Fig. 3C, D, left). This ramp was less apparent for D2+ neurons (Fig. 3C, D, right), whose firing rarely distinguished engaged vs. non-engaged trials.

**Fig. 3.**
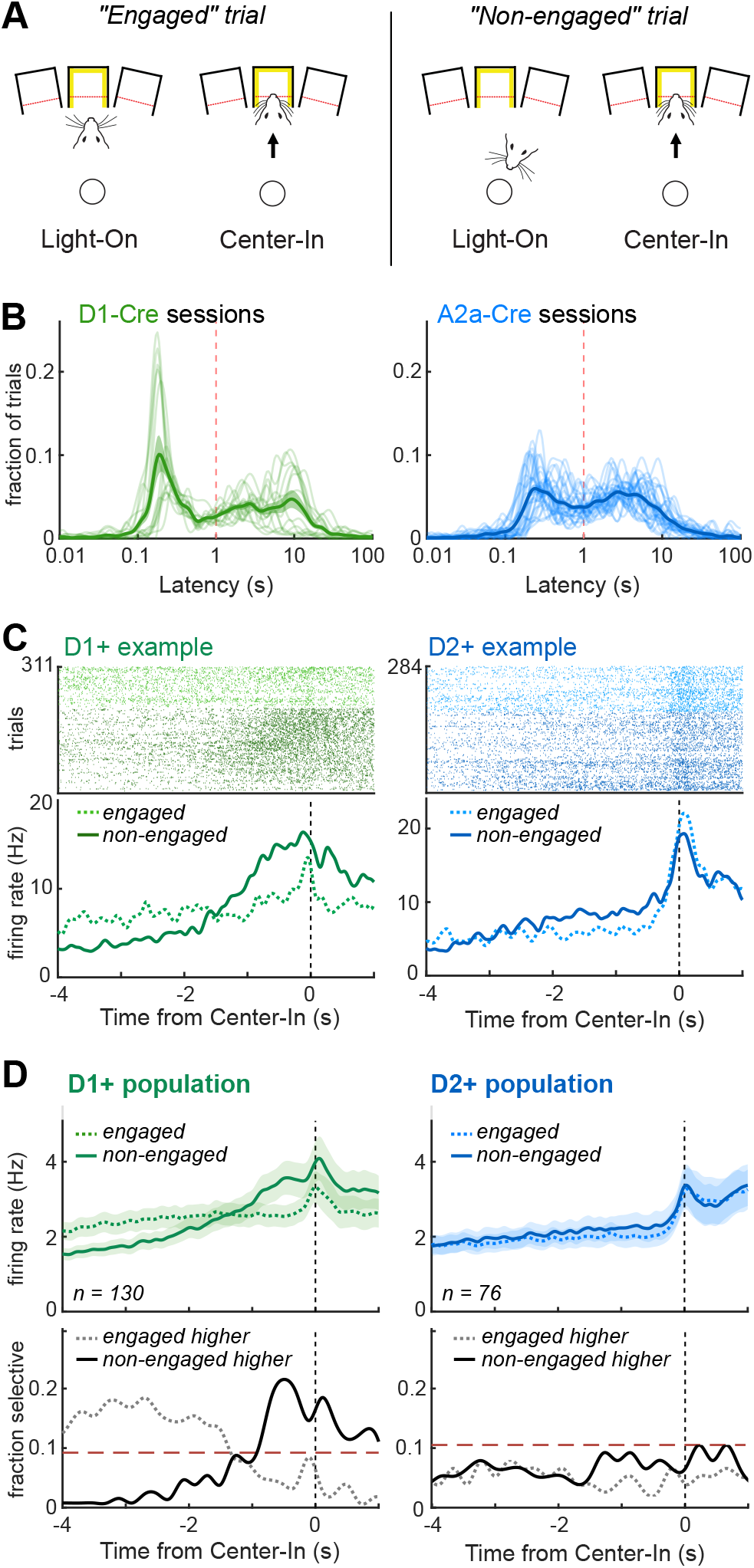
D1+, but not D2+, SPNs are preferentially active during motivated approach. **A**, Left, on “engaged” trials the rat is already waiting at the center port for the Light-On event, and as a result latencies are very short. Right, for “non-engaged” trials the rat is at various locations in the chamber at Light-On, and the time required to decide to work and approach the center port produces longer latencies. **B**, Bimodal latency distributions for both D1-Cre and A2a-Cre rat sessions. Engaged and non-engaged are classified by the red boundary at 1 s. **C**, Examples of individual D1+ (left, green) and D2+ (right, blue) SPN neuron activity. Top, rasters of spikes on each trial; bottom, histogram of firing rate (50 ms gaussian kernel convolution), each aligned on Center-In. **D**, Subpopulation activity, shown as average firing (top) or fractions of cells firing significantly more for engaged compared to non-engaged trials, or vice-versa (bottom, permutation tests, p < 0.05, Bonferroni corrected). Dashed lines indicate when fractions are significantly higher than chance (binomial threshold, p < 0.05).

Rats are more likely to decide to approach the Center port when reward rate (trial value) is high (Supplementary Figure 2). Correspondingly, we found that the approach-related ramp in D1+ firing is stronger when trial value is higher (Fig. 4A, B). A significant fraction of D1+ SPNs showed this form of positive value coding during approach (Fig. 4B, bottom left), with no indication of the opposite, negative value coding (more firing when reward is less expected). The D2+ SPN population showed neither positive nor negative value coding during motivated approach (Fig. 4B, bottom right). D1+ value coding was much less apparent in engaged trials (Supplementary Fig. 3A), for which no substantial approach is needed.

**Fig. 4.**
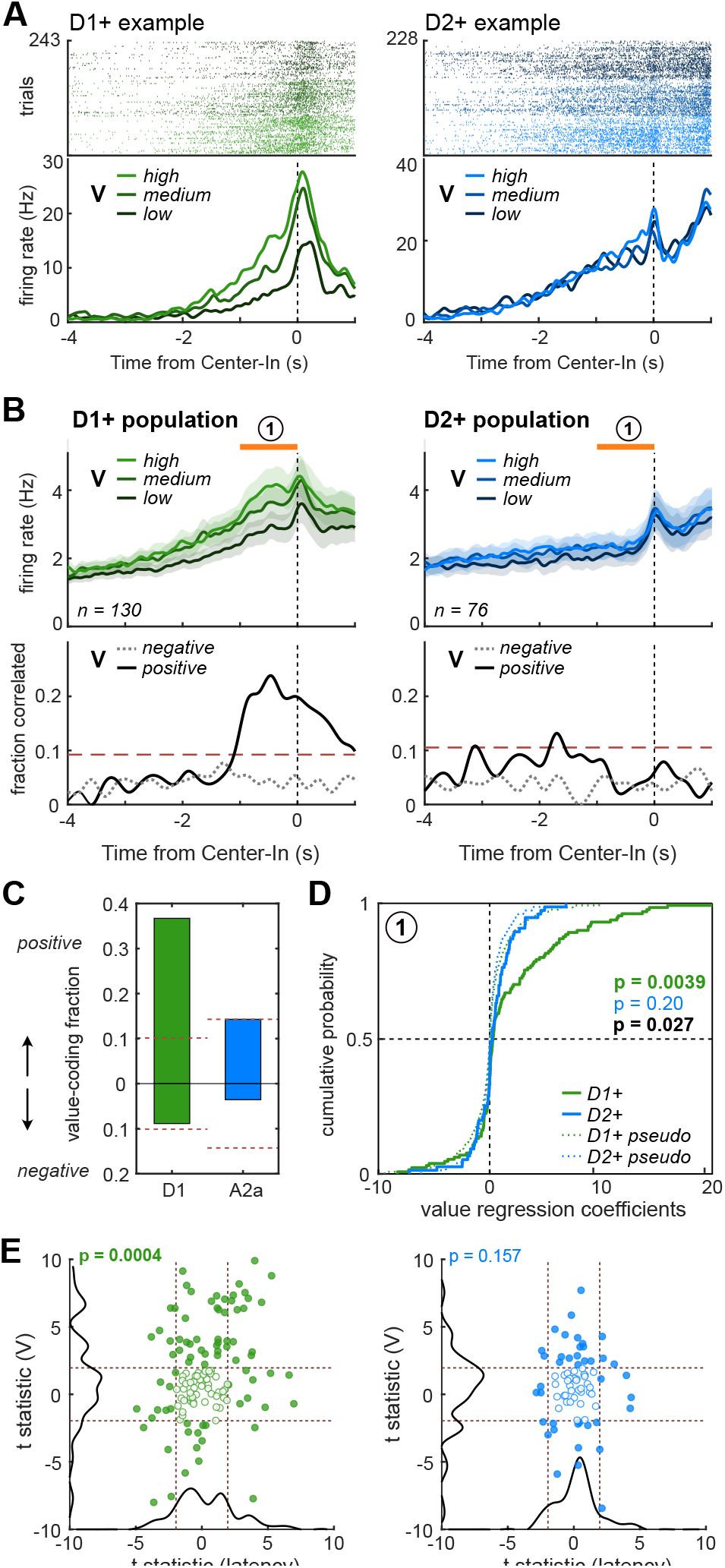
D1+, but not D2+, SPNs signal value during approach. **A**, Examples of individual D1+ (left, green) and D2+ (right, blue) SPN neuron activity during non-engaged trials only, split into terciles of trial value (low, medium, high). Both of these example cells ramp up during approach, but only the D1+ firing depends on reward expectation. **B**, Subpopulation activity, shown as average firing (top) or fractions of cells showing significant positive or negative correlations with trial value (p < 0.05, Pearson’s correlation, Bonferroni corrected). Dashed lines indicate when fractions are significantly higher than chance (binomial threshold, p < 0.05). **C**, Fractions of active D1+ and D2+ units having significant positive correlations (positive fraction) or negative correlations (negative fraction) during the 1 s prior to Center-In on non-engaged trials (orange bar “1” in B, used for all analyses in panels C-E), corrected for spurious correlations. **D**, Cumulative distributions of regression coefficients for trial value during approach. Shown p values are from Kolmogorov-Smirnov tests between D1+ and D2+ distributions (black), D1+ and average pseudo-session distribution (green), and D2+ and average pseudo-session distribution (blue). **E**, Multiple regression analyses incorporating both trial value (V) and latency. Plotted are t statistics for D1+ (left, green) and D2+ (right, blue) cell responses. p values indicate that trial value contributes significantly more to D1+ firing than latency, but this is not the case for D2+ cells (permutation tests between t statistics). Dashed lines indicate thresholds for significant regression (t>1.96, t<-1.96).

We obtained similar results if we considered the total spiking of each cell in the 1s epoch before Center-In (Fig. 4C-E). After correcting for spurious correlations (Harris 2020), 35.2 % of D1+ SPNs but only 10.7% of D2+ neurons showed positive value coding during approach (Fig. 4C; significantly different fractions, p = 0.0146, Fisher’s exact test). For D1+ cells, the distribution of regression coefficients for value coding was significantly different to chance (using pseudo-sessions to generate a null distribution, see Methods), and also different to the D2+ population, which itself was not significantly different to chance (Fig. 4D). This D1+ result was unchanged if we estimated value in other ways, such as with a leaky integrator or Q-learning (Supplementary Fig. 3B), and was also apparent in the data from each of the D1-Cre rats individually (Supplementary Fig. 3C). To gain further confidence that this value coding result did not arise from differences in overt behavior, we performed a multiple regression analysis on each neuron’s firing rate, incorporating both trial value and latency. For D1+ cells, value coefficients were significantly stronger than latency coefficients (Fig. 4E, left); this was not the case for D2+ cells (Fig. 4E, right). We conclude that a subset of D1+ SPNs show a selective, reward-expectation-dependent increase in firing during motivated approach.

### Value-modulated increases in D1+ and D2+ firing during choice execution

We then assessed how value-modulation of activity evolves through the remainder of the trial (illustrated for all cells in Supplementary Figure 4A). We considered the subset of neurons with increased firing during the hold period before the Go cue, and/or the subsequent withdrawal and lateral movement before Side-In (Fig. 5). The D1+, but not D2+, subpopulation showed positive value coding during the hold period (Fig. 5A,B bottom left panels). Subsequently, as rats performed their leftward/rightward movement (between Center-Out and Side-In) both D1+ and D2+ cells increased average firing (more prominently for D2+), and both cell types showed an increase in positive value coding (Fig. 5A,B). This observation of higher reward expectation selectively elevating D1+ firing during the hold period, then elevating both D1+ and D2+ firing during the left/right movement, was also visible in the distributions of regression coefficients (Fig. 5C). However this did not reach significance for D2+, possibly due to fewer cells included in the analysis.

**Fig. 5.**
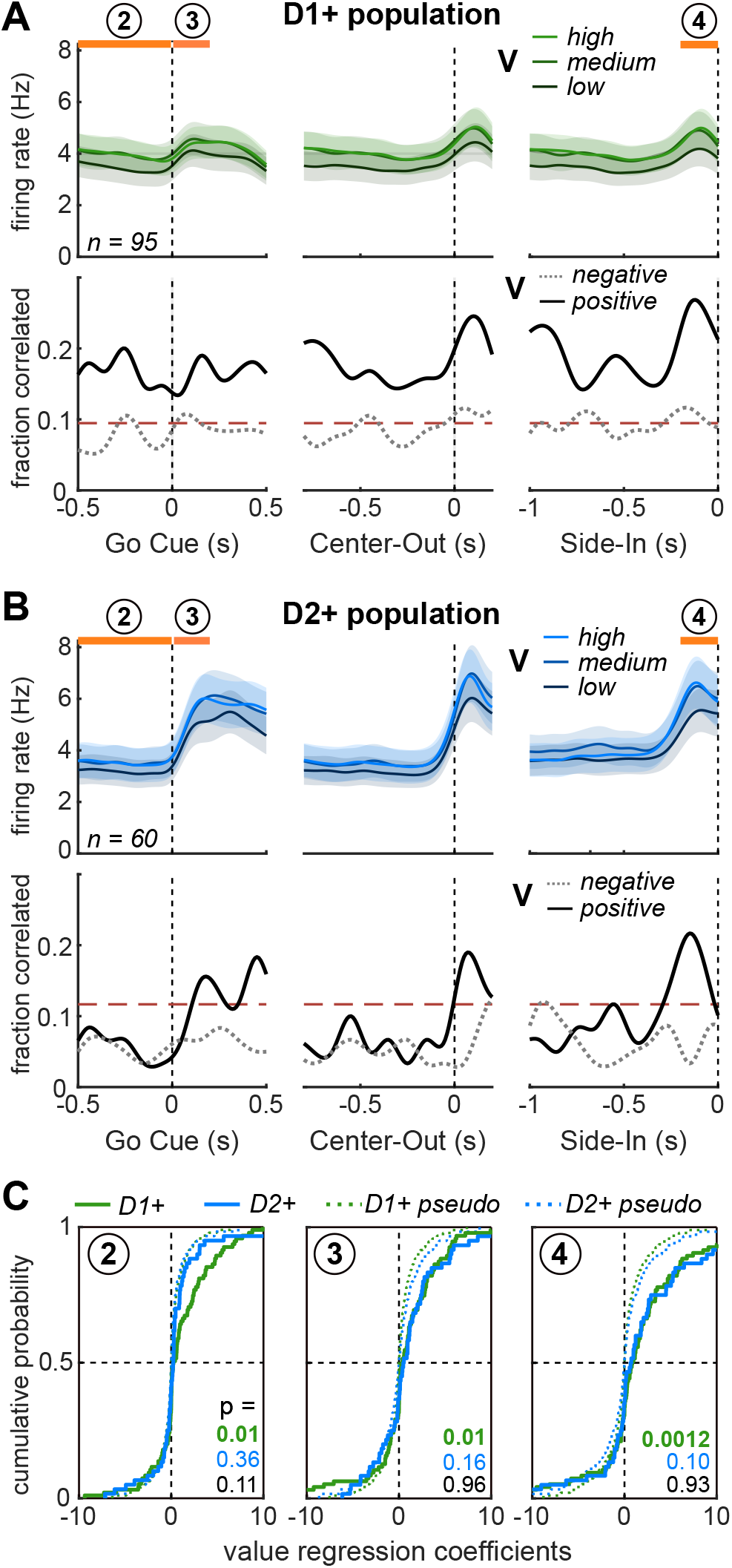
Value coding by D1+ and D2+ SPNs during choice execution. **A**, Top, mean firing of D1+ SPNs from the last part of the hold period until Side-In, split into terciles of trial value (V). Neurons included in this figure are those with average firing rate increases during any of the histogram ranges shown (i.e. 0.5s before Go cue to 0.5s after; 0.8s before Center-Out to 0.2s after; 1s before Side-In until Side-In). Bottom, fraction of D1+ SPNs at each moment whose firing has significant positive (V+) or negative (V-) correlations with value (p < 0.05, Pearson’s correlation, Bonferroni corrected). Dashed lines indicate when fractions are significantly higher than chance (binomial threshold, p < 0.05). Orange labelled bars indicate time epochs used for analysis in C. **B**, Same as A, but for D2+ SPNs. **C**, Cumulative distributions of regression coefficients, for the dependence of SPN firing on trial value V. Regression analysis was performed separately for each of the numbered epochs in A-D. p values shown are from Kolmogorov-Smirnov tests comparing D1+ to average pseudo-session distribution (green); D2+ to average pseudo-session distribution (blue); D1+ to D2+ groups (black).

Subsets of both D1+ and D2+ cells distinguished between left and right movements, with a preference for the contraversive direction (i.e. moving towards the opposite side from the brain hemisphere in which the neuron was recorded; Supplementary Fig. 5). However, this coding of movement direction was sparse and weak compared to trial value coding during movement (Supplementary Fig. 4). We also examined whether value coding by NAc cells might reflect not just the history of rewards vs omissions, but whether specific left/right choices had been recently rewarded. Such “action-value” coding has been previously reported for striatal neurons (28, 44, 45), including in accumbens (46). We employed a trial-based Actor-Critic model (see Methods) that tracks both trial value as before (via the Critic component) and also reward-history-dependent preferences for left and right choices (the actor component “weights”). We regressed each neuron’s firing during movement (epoch “4”) against each of these decision variables. Both D1+ and D2+ neurons showed stronger relationships to trial value than to action weights (for ipsiversive, contraversive, or the chosen direction; permutation tests on t statistics; Supplementary Fig. 6).

**Fig. 6.**
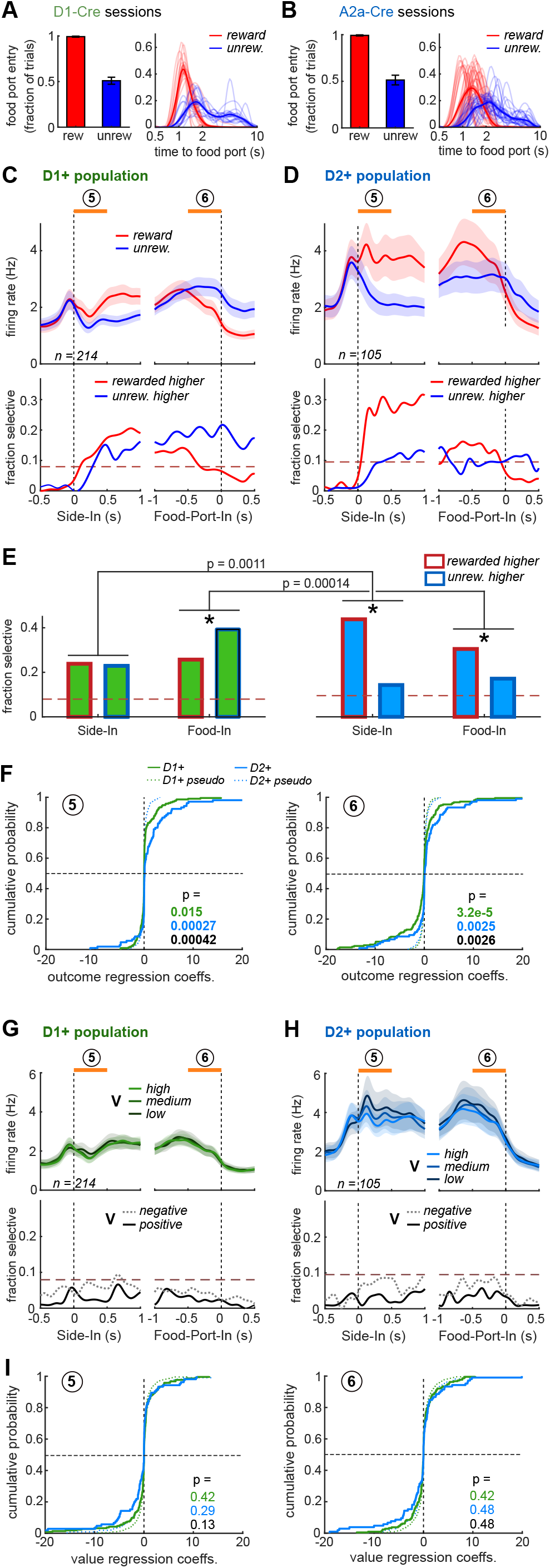
Divergent outcome responses in D1+ and D2+ SPNs. **A**, Left, Fraction of rewarded and unrewarded trials during which D1-Cre rats approach and enter the food port. Right, distributions of time duration of D1-Cre rats to enter food port on rewarded and unrewarded trials (faint lines – individual sessions, bold lines – average distribution mean +/-SEM). **B**, Same as A, but for A2a-Cre rats. **C-D**, Top panels, mean firing of task-related D1+ and D2+ cells at Side-In and food port entry, split by trial outcome. Orange labelled bars indicate time epochs used for analysis in E-F. Bottom panels, fraction of units selective for rewarded vs. unrewarded trial outcomes (permutations tests, p < 0.05, Bonferroni corrected). Dashed lines indicate when fractions are significantly higher than chance (binomial threshold, p < 0.05). **E**, Fractions of D1+ and D2+ units selective for outcome type following outcome reveal and during food port approach (500 ms after Side-In 500 ms before Food-In, epochs “5” “6” in C-D). * p < 0.05, chi-square distribution. **F**, Distributions of regression coefficients for outcome (rewarded or unrewarded) for D1+ and D2+ cells following outcome reveal (5) and during food port approach (6). p values shown are from Kolmogorov-Smirnov tests between D1+ and average pseudo-session distribution (green), and D2+ and average pseudo-session distribution (blue), D1+ and D2+ groups (black). **G-H**, Top panels, mean firing of task-related D1+ and D2+ cells at Side-In and food port entry, split into terciles of trial value (V) for rewarded outcomes. Bottom, fraction of D1+ SPNs at each moment whose firing has significant positive (V+) or negative (V-) correlations with value (p < 0.05, Pearson’s correlation, Bonferroni-corrected). Dashed lines indicate significance thresholds (binomial tests, p < 0.05). Orange labelled bars indicate time epochs used for analysis in I. **I**, Cumulative distributions of regression coefficients, for the dependence of SPN firing on trial value V following outcome (5) and during food port approach (6). p values shown are as in F.

### Reward delivery preferentially activates NAc D2+ neurons

At the completion of the leftward/rightward movement (Side-In), rats may or may not hear the reward cue (food delivery click). If they do, they leave the side port and approach and enter the food port, promptly and consistently (>99% of trials; Fig. 5A,B). Many NAc D1+ and D2+ neurons increased firing during this overall period (from 0.5s before Side-In to 0.5s after Food-Port-In; Fig. 5C, D). However, the reward cue was much more likely to elicit increased firing of D2+, compared to D1+ cells (Fig. 6D-F, epoch “5”; Supplementary Fig. 2). This was surprising as D2+ cells are generally thought to respond to disappointing, no-reward events (31, 32, 47). As rats approached the food port, there was a preferential increase in D1+ cells specifically on trials in which food was not delivered (Fig. 6C-F, epoch “6”).

In contrast to earlier portions of the trial, neither D1+ nor D2+ populations showed positive value coding after Side-In (Fig. 6G-H). Both cell types appeared to show a greater degree of negative value coding shortly after hearing the reward cue (Fig. 6G-H, epoch “5”), and this pattern was clear for some individual D2+ cells (Supplementary Fig. 7). Such negative modulation of reward cue responses by reward history would be consistent with RPE coding (as is seen for NAc DA release in this task; (17)). However, the proportion of each subpopulation exhibiting this negative value coding did not cross a strict statistical threshold. Similar results were seen for unrewarded trials (Supplementary Fig. 8). We conclude that such RPE coding may be present in NAc, but at most in sparse subsets of each cell type ((48); Supplementary Fig. 4).

## Discussion

### NAc Core D1+ cells and value-modulated approach

We have shown that as rats begin a trial by approaching a port, NAc Core D1+ SPNs ramp up their firing in proportion to the rats’ expectation of being rewarded later in the trial. Though not - to our knowledge - demonstrated before, this result fits well with prior studies. The NAc Core is closely involved in motivation to work to obtain rewards (“appetitive” or “incentive” behavior;(49–52)). Subsets of (unidentified) NAc cells have been previously reported to increase spiking in apparent anticipation of rewards (25), after reward-predictive cues that evoke approach responses (53–55), and in mazes as animals run towards rewards (56). Furthermore, imaging studies have reported selective increases in D1+ calcium transients as animals approach places that are preferred due to prior pairing with cocaine (57) or heroin (58).

Of note, there was no comparable increase in NAc D1+ firing after the reward cue at Side-In, rats moved from the side port to the food port to collect delivered rewards. The NAc Core is not typically needed when cues instruct fixed actions, in subjects already engaged in a task. Rather, it is key for motivating “flexible” approach behavior (6)-e.g. when starting a trial from a variable location. Experimentally, one way to generate the need for flexible approach is to use long intertrial-intervals, so that rats disengage from task performance between trials (26). Under these conditions, cues that reliably signal reward availability can evoke robust and rapid increases in NAc Core firing, accompanied by consistent, prompt approach behavior (59). Instead of using long intervals, our task varies the probability that work will be rewarded. As a result, in non-engaged trials the approach behavior is largely self-paced. The Light-On cue elevates the hazard rate (instantaneous likelihood) of initiating approach (33), but latencies remain highly variable. Under these conditions, we found only modest NAc firing to the Light-On cue, and instead a ramping increase in NAc D1+ firing accompanying the approach. This approach-linked activity was greater when rats had a higher expectation of reward. Critically, this reward expectation was based on internal tracking of reward history, rather than external cues that explicitly signal reward probability or magnitude. These observations provide important new evidence for the role of NAc D1+ cells in motivated decisions that are internally generated, rather than elicited by the “incentive salience” of discrete external stimuli (60).

### Influence of DA over D1+ and D2+ cells

Decisions to work depend critically upon DA modulation of the NAc (7). Blockade of NAc Core dopamine receptors - especially D1r-interferes with the initiation of approach behavior (34, 61, 62). Mice lacking functional D1r show a deficit in cue-evoked approach, and this deficit is ameliorated by restoring expression of D1r specifically in NAc Core D1+ neurons (63). Conversely, many stimulant drugs achieve their psychomotor activating effects through increased NAc DA release and binding to D1r (64–66). The critical time scales over which DA modulates behavior are an active area of discussion (8, 67–69). In slice studies, activation of D1r can enhance the excitability of D1+ neurons within hundreds of ms (10). Furthermore, the fast (subsecond-second) dynamics of NAc DA release show some striking relationships to the D1+ activity we report here. Like D1+ firing, DA release has also been shown to ramp up during motivated approach (70**?**), with greater ramping when reward expectation is higher (71, 72). In our specific task, we previously observed a fast ramp in DA release before Center-In (17). Furthermore, selective optogenetic stimulation of VTA DA cells enhanced the hazard rate of approach immediately (again, within a few hundred ms; (33)). This convergent evidence suggests that fast DA dynamics are rapidly shaping D1+ excitability based on reward expectation, to influence decisions to approach. However, we are not directly demonstrating this here. Moreover, at least in slices D1r-enhanced excitability may not reset quickly enough to enable D1+ cells to track fast fluctuations in DA release (10).

D1r and D2r affect SPN excitability in opposite ways (11, 73, 74). We might therefore expect that approach-related increases in DA would be accompanied by a ramp downwards in D2+ firing, which was not apparent. One possibility is that D2r typically exists in a higher affinity state (75) such that DA present before approach is already enough to saturate D2r. If true, D2r would be only responsive to decreases in DA, and the DA approach ramp would be undetectable by D2+ cells. However, other evidence from striatal brain slices indicates that D2r are functionally low-affinity, and D2+ cells can rapidly respond to DA increases (76). In awake behaving animals the affinity of DA receptors is unknown, and there is also great uncertainty about “baseline” DA concentrations (8, 77, 78). These represent substantial barriers to understanding striatal circuit dynamics that should be addressed in future work.

Furthermore, the patterns of D1+ and D2+ firing after reward delivery or omission did not obviously reflect rapid modulation by DA. In particular, the dip in DA that occurs with omission (17) would be expected to enhance D2+ excitability, and thus firing. Others have reported such a pattern (30), notably in dorsomedial striatum (31, 32). However we found that few NAc D2+ cells fired more to omissions than reward delivery - instead, a much larger proportion showed the opposite pattern, increasing firing at a time we previously reported a strong pulse of DA release. We also expected that this DA pulse would be accompanied by greater D1+ firing, especially as artificial stimulation of NAc D1+ cells can be strongly reinforcing (79). Instead similar numbers of D1+ preferred reward delivery or omission. Of course, extracellular recording of spikes provides only limited information about intracellular processes, and spiking also depends on complex patterns of glutamate and GABA input, among others. Nonetheless, studies of accumbens synaptic plasticity (20) and models of the resulting reward-related learning (22, 47, 80) generally emphasize the importance of strong post-synaptic depolarization, as part of a three-factor plasticity rule (presynaptic, postsynaptic activity, plus dopamine modulation; (81–83)). Our results suggest a need for updated models of striatal learning mechanisms, particularly in NAc.

### Separate and common patterns of D1+ / D2+ firing

Despite long-standing evidence that D1+ and D2+ SPNs have opposing functional roles, over the last decade several groups have reported that both cell types jointly increase activity during movements (84–86). These activity increases are preferentially seen for contraversive movements, and manipulations of either cell type interfere with such contraversive movements (87). In NAc we also found that both D1+ and D2+ have movement-selective activity with a contraversive bias. However, movement-selective firing was sparse in NAc (Supplementary Fig. 4), consistent with earlier observations (88). This presumably reflects the lower level of NAc involvement in lateralized orienting decisions compared to other subregions, especially dorsomedial striatum (caudate; (89–91)). Future studies from our laboratory will examine and compare dorsal striatal D1+ and D2+ activity in the same task used here.

During this lateralized movement both NAc D1+ and D2+ fired more when more recent trials had been rewarded. Reward expectation has long been reported to modulate striatal neurons, including cells involved in orienting-type movements (92). We found NAc SPNs were modulated by overall reward history, more than the history of rewards for specific left/right choices (44). This is also consistent with a range of prior results for ventral striatum (46, 88, 93, 94) and the general idea that NAc is concerned with values of states, rather than values of actions (95). It has been repeatedly proposed that movement-related cells positively modulated by value are D1+, and those negatively modulated are D2+ (e.g. (22, 96)). Our results do not support this hypothesis, at least for NAc (see also (97)). It is possible that both D1+ and D2+ value coding during movement execution simply reflects information already present in their glutamatergic inputs. Both SPN cell types in NAc Core receive strong input from structures that are themselves value-modulated (Li et al. 2020), including medial frontal cortex ((17, 98)) and basolateral amygdala ((55, 99)). As a result, certain aspects of value coding may be better thought of as an overall network state (100) rather than a readout of learned corticostriatal synaptic weights. Regardless of how it arises, such value coding - present immediately before animals receive feedback about reward at Side-In - could contribute to the downstream computation of reward prediction errors. D1+ provide a major input to DA neurons (e.g. (101)), while D2+ project to globus pallidus, where we have recently described a population of cells that encode RPE in a remarkably similar fashion to DA neurons (102).

### Contributions of NAc D2+ neurons

At other times in the task, the activity patterns of D1+ and D2+ populations were highly distinct. In contrast to the selective D1+ increase during approach, D2+ cells preferentially increased activity after the Go cue until entry into the food port. The function of this D2+ population change is currently unclear. D2+ cells have been previously argued to suppress alternative actions to that selected (103, 104). Following this idea, the NAc D2+ increase seen here may help maintain ongoing trial performance, helping the rat stay motivated instead of deviating into a different course of action (23). This may relate to observations that lower NAc D2r expression is correlated with impulsivity, in other tasks (105). It is not obvious, however, why such D2+ activity would begin at the Go cue rather than during the preceding hold period. Furthermore, as noted by others (e.g. (12)) the fact that D2+ cells show similarly detailed and varied patterns of firing to D1+ neurons argues against their providing a simple “suppress everything else” signal. Developing insightful new tests of NAc D2+ functions is an important goal for future research.

## ACKNOWLEDGEMENTS

We thank Howard Fields and Wei Wei for providing valuable feedback on manuscript drafts, and Lily Pelattini and Kevin Kyoungjun Kim for technical assistance. This study was supported by the National Institute on Drug Abuse, the National Institute of Neurological Disorders and Stroke, the State of California, and the University of California, San Francisco.

## AUTHOR CONTRIBUTIONS

TWF performed electrophysiological recordings and data analyses. AM contributed code for analyses and reinforcement learning models. JDB designed and supervised the study and obtained funding. The manuscript was written by TWF and JDB.

## Methods

### Animals

All animal procedures in this study were approved by the University of California San Francisco Institutional Committee on Use and Care of Animals. Adult rats (300–500 g; all males as their greater size made them better able to tolerate the large implant), hemizygous D1-Cre or A2a-Cre on a Long-Evans background (35) were maintained on a reverse 12:12 light:dark cycle and tested during the dark phase. Rats were mildly food deprived (>85% free weight), receiving 15 g of standard laboratory rat chow daily in addition to food rewards earned during task performance.

### Bandit task

Pretraining and testing were performed in computer-controlled Med Associates operant chambers (25 cm × 30 cm at widest point) each with a five-hole nose-poke wall, as previously described (33). Rats were shaped and trained for the bandit task over 3+ months, and after implantation were recorded for 1-2 months. Recordings took place every other day to allow for tetrode locations to stabilize and rat motivation to perform the task to remain high.

### Electrophysiology

Rats were bilaterally implanted with custom-designed drivable tetrode arrays, each consisting of 16 tetrodes (constructed from 12.5-μm nichrome wire, Sandvik) mounted into a circular pattern (652 μm diameter) around a stationary central tapered optic fiber (200 μm, Optogenix). During the same surgery, we injected 1 μl AAV5-Syn-FLEX-rc[ChrimsonR-tdTomato] into nucleus accumbens core (AP +1.75 mm, ML +/-1.6 mm, DV −6.5 or-7.0 mm). Optic fiber initial depths were 6.3 or 6.5 mm, tetrode initial depths were 5.8 or 6.0 mm, below brain surface. Wideband (1–9,000 Hz) brain signals were sampled (30,000 samples / s) using custom headstages built around Intan amplifier chips. Optrodes were lowered 80 μm at the end of each recording session, so that the same cells were not repeatedly recorded. Individual units were isolated offline using a MATLAB implementation of MountainSort (106) followed by careful manual inspection.

#### Classification

Units were classified as putative SPNs based on waveform and firing rate properties (peak-valley duration >560 μs, peak half-width duration >100 μs, session-wide firing rate <10 Hz). Light-responsive or tagged units were identified using the stimulus-associated latency test (Kvitsiani et al. 2013). In brief, at the end of each experimental session, we connected the optic fiber to a laser diode and delivered light pulse trains of 100 pulses (10 ms) at 1 Hz, using 1, 3, 10, or 20 mW light power calculated for the fiber tip. For a unit to be identified as light-responsive it needed to reach the significance level of P = 0.001 for 10-ms pulse trains and respond during the 10 ms light pulse on at least 3% of pulses in a train, for at least one power level. We also compared the light evoked waveforms (within 10 ms of laser pulse onset) to session-wide averages; all light-responsive units included in the data set had a Pearson correlation coefficient > 0.9. D1+ neurons were successfully recorded from three D1-Cre rats (IM-1136, 31 units; IM-1371, 295 units; IM-1399, 172 units) over 14 sessions. D2a+ neurons were recorded from four A2a-Cre rats (IM-1154, 39 units; IM-1199, 24 units; IM-1373, 95 units; IM-1388, 106 units) over 29 sessions. Three D1-Cre rats were excluded from this dataset due to recording locations entirely outside of the nucleus accumbens Core. A total of 5 additional rats (2 D1-Cre, 3 A2a-Cre) were implanted with other drive designs and failed to produce any tagged units.

### Modeling

The primary reinforcement learning model throughout this work is a simple trial-based “Critic”. This tracks a single quantity: trial value, V. On each trial the outcome (reward = one, no reward = zero) is used to calculate a reward prediction error, RPE = outcome - V. The trial value is then updated for the next trial using V’ = V + alphaC * RPE. AlphaC is the Critic learning rate, which was fit to behavior, maximizing the relationship between V and (negative) log (latency). For all behavioral fits, we first determined the best-fit parameter for each session individually, then applied of this parameter distribution to all sessions.

In supplementary figures we also used some other simple models, as follows. The leaky integrator model (used as before, (33)) again tracks a single decision variable, reward rate. This increases by one at each reward receipt, and decays exponentially in time. The decay parameter was chosen to maximize the relationship between reward rate and (negative) log (latency). The trial-based Actor-Critic model used the same Critic as before, but the additional Actor tracked action weights for leftward and rightward choices. For the action chosen on each trial, the weight was updated using the same RPE as for the Critic, with W’(chosen) = W(chosen) + alphaA * RPE. For the Q-learning model there was no Critic, instead for each action (left, right) the model tracked an action value Q. This was updated for the chosen action using Q’(chosen) = Q + alpha*RPE, with the RPE here being the difference between the outcome and the Q value of the chosen port. The Actor learning rate alphaA and the Q-learning rate alpha were both fit to maximize the relationship between policy and choice. The overall trial value for the Q-learning model was estimated as the sum of both Q values (Qleft + Qright).

### Permutation tests

To determine whether units had significantly increased or decreased activity around an event of interest, we shuffled trial timestamps pseudo-randomly throughout the bandit task session and averaged the firing rate of the shuffled timestamps (200 ms bin size). This was repeated 10,000 times. A unit’s average firing rates around an event of interest were considered significant if they were above or below the permutated distribution (p<0.05, two-tailed, corrected for multiple comparisons). To test whether units significantly differentiated between two test conditions (engaged vs. unengaged; ipsiversive vs. contraversive; rewarded vs. unrewarded), we shuffled the trial IDs 10,000 times and created a permutated distribution of firing rate differences between these two shuffled conditions. If the actual difference in firing rates between the two conditions was significantly above or below the permutated distribution (p<0.01, two-tailed, corrected for multiple comparisons), the unit was considered to be selective for that test condition.

### Correction for spurious correlations

In tasks comparing activity to decision variables such as value, spurious correlations can arise due to temporal dependencies (i.e. adjacent trials should not be treated as entirely independent; (107)). These were corrected for using the pseudo-session method (Harris 2020). Briefly, we assembled a collection of bandit task sessions from multiple Berke Laboratory experiments, choosing sessions with >150 trials, >50% of initiated trials successfully completed, and <80% choice for each side-port. If trial numbers were shorter than the real trial numbers for a given bandit task session, then two pseudo-sessions were randomly concatenated to match the previous block probabilities at the end of one pseudo-session and the continuation on the next. This produced 5,098 pseudo-sessions with which to test whether correlations were spurious. A correlation was considered spurious if the real session correlation coefficient failed to exceed the distribution of pseudo-session correlation coefficients (p<0.05, two-tailed).

**Extended Data Figure 1.**
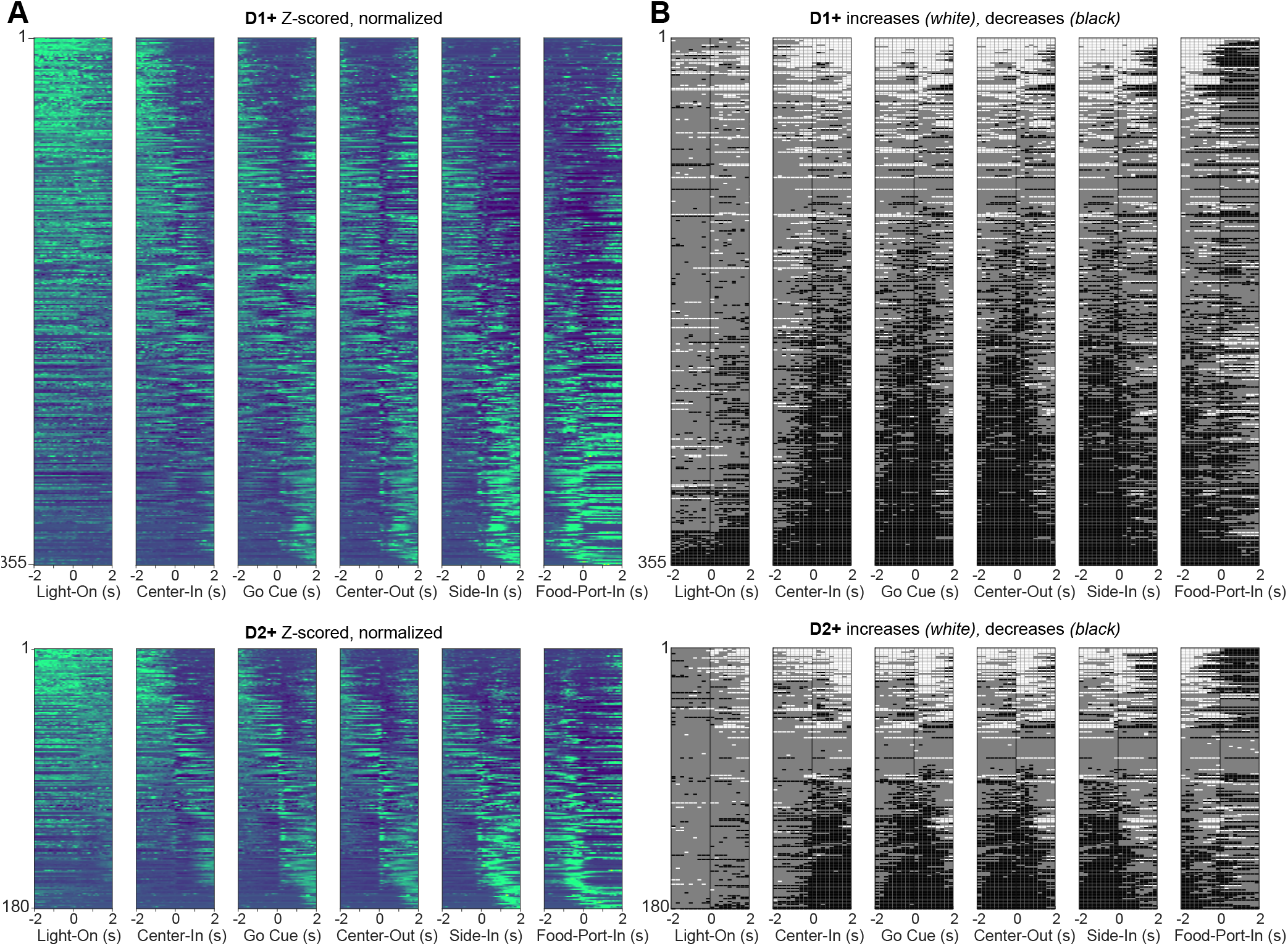
Firing patterns of all task-related D1+ and D2+ neurons. **A**, Z-scored, normalized firing for each unit in each subpopulation. Color scale is blue (minimum) to green (maximum), with colors saturated to make dynamics more visible. Cells are sorted by center of mass. **B**, Temporal pattern of firing increases and decreases in firing for each task-related D1+ and D2+ neuron. White (black) bars indicate significant increases (decreases), compared to permutated distribution (p < 0.05, Bonferroni corrected). Cells are sorted by overall extent of modulation (positive – negative). All data after Side-In are from rewarded trials only.

**Extended Data Figure 2.**
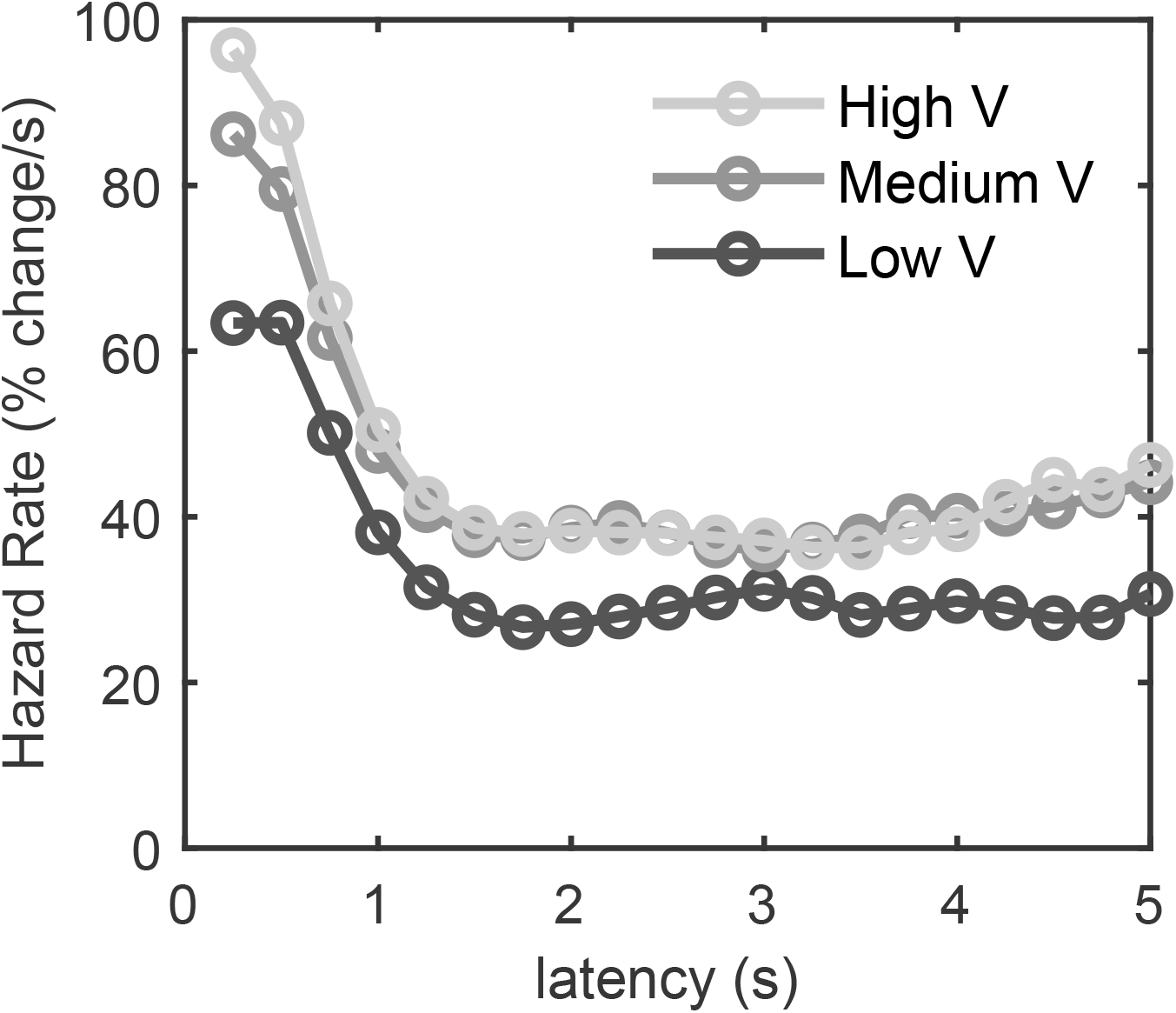
Hazard rate of trial initiation by trial value. Probability that trials will be initiated if they have not already been initiated, split into terciles of trial value (V).

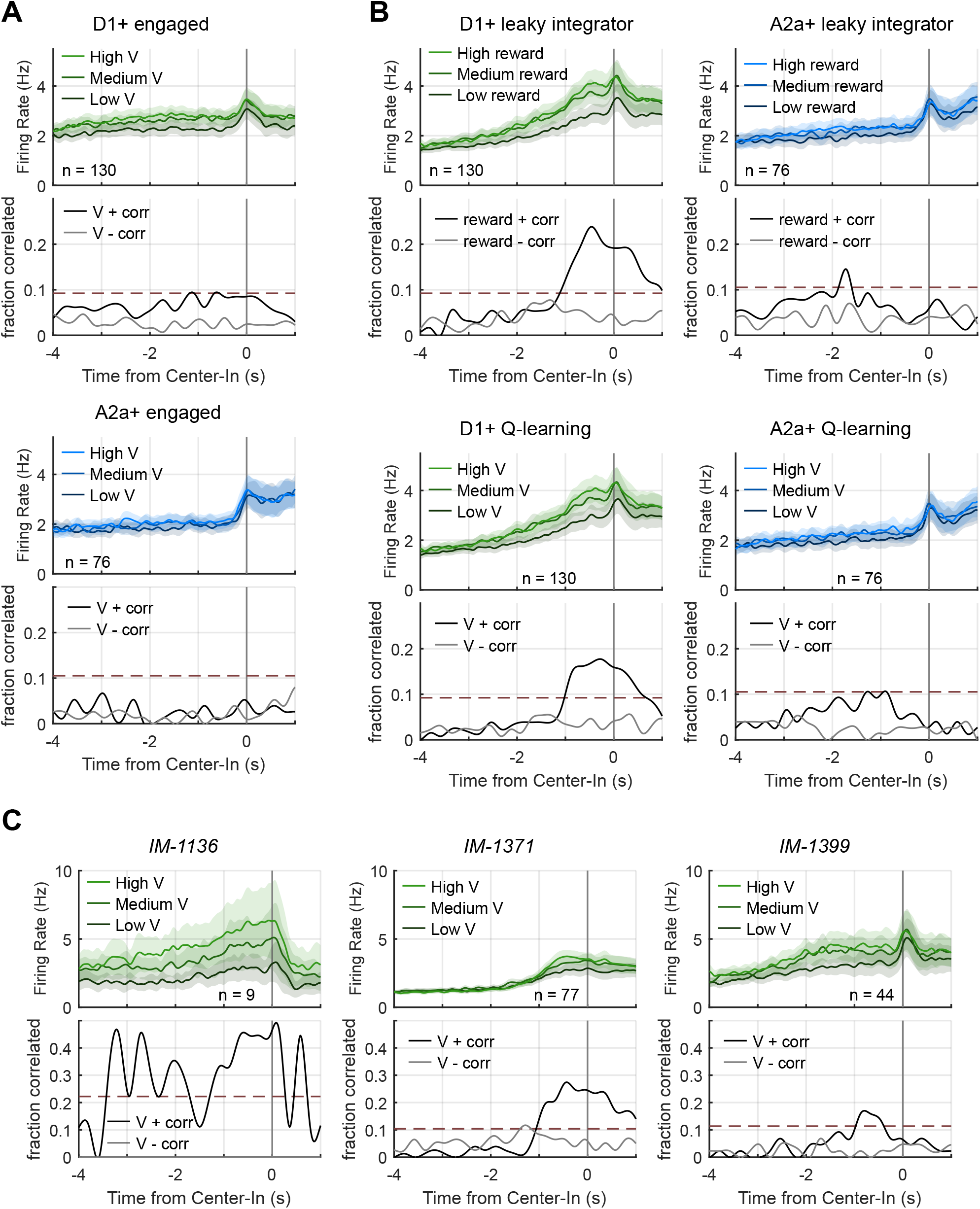

**Extended Data Figure 4.**
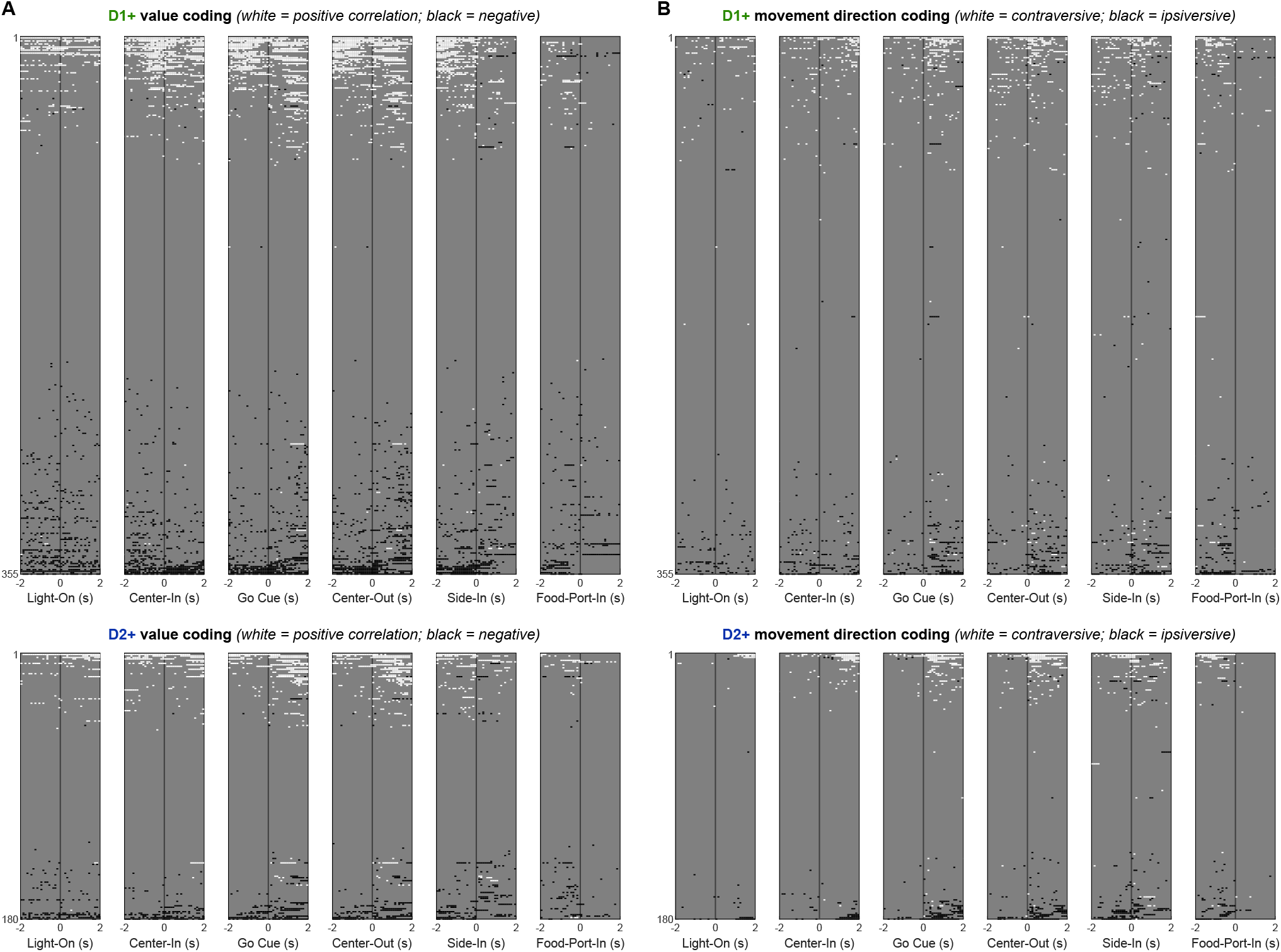
Significant value and choice responses of all task-active D1+ and D2+ neurons during the bandit task. **A**, Value correlations of all task-active D1+ and D2+ neurons. White bars indicate positive correlations and black bars indicate negative modulation (p < 0.05, Bonferroni corrected). **B**, Choice responses of all task-active D1+ and D2+ neurons. White bars indicate contraversive and black bars ipsiversive selectivity (permutation test, p < 0.05, Bonferroni corrected). Cells are sorted as in Supp. Fig. 1B. All data after Side-In are from rewarded trials.

**Extended Data Figure 5.**
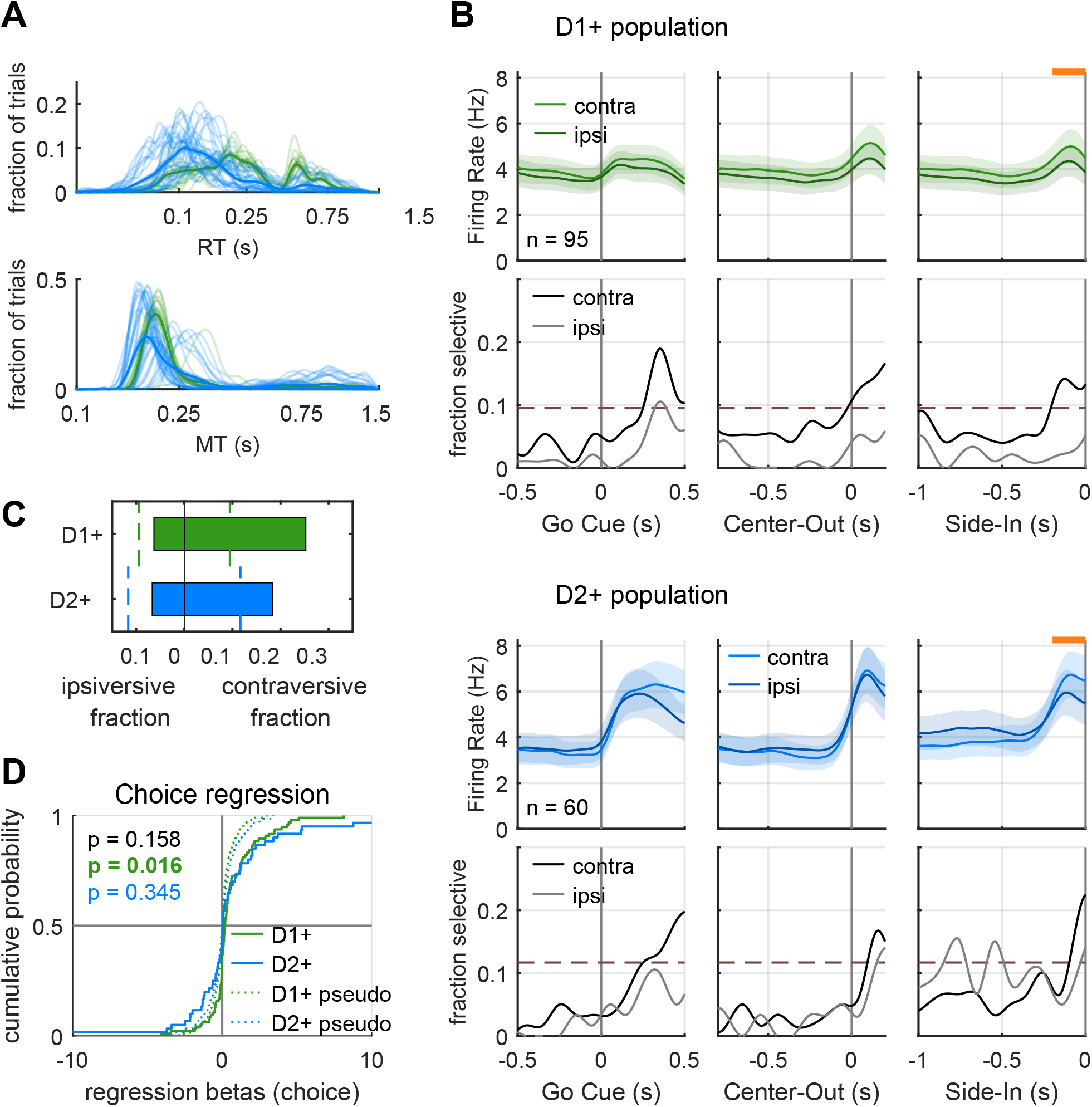
Both D1+ and D2+ cells are preferentially active for contraversive actions. **A**, Reaction time (top, Go Cue to Center-out) and movement time (bottom, Center-out to Side-In) distributions for D1-Cre and A2a-Cre rats, as in Fig. 3B. **B**, Contraversive and ipsiversive responses of D1+ and D2+ cells as in Fig. 5A-B (permutation test, p < 0.05, Bonferroni corrected). **C**, Fraction of D1+ and D2+ cells selective for choice direction during choice (200 ms prior to Side-In, See Fig. 5). **D**, Distributions of choice regression coefficients for D1+ and D2+ cells during choice. p values shown are for Kolmogorov-Smirnov tests between D1+ and average pseudo-session distribution (green), and D2+ and average pseudo-session distribution (blue), D1+ and D2+ groups (black).

**Extended Data Figure 6.**
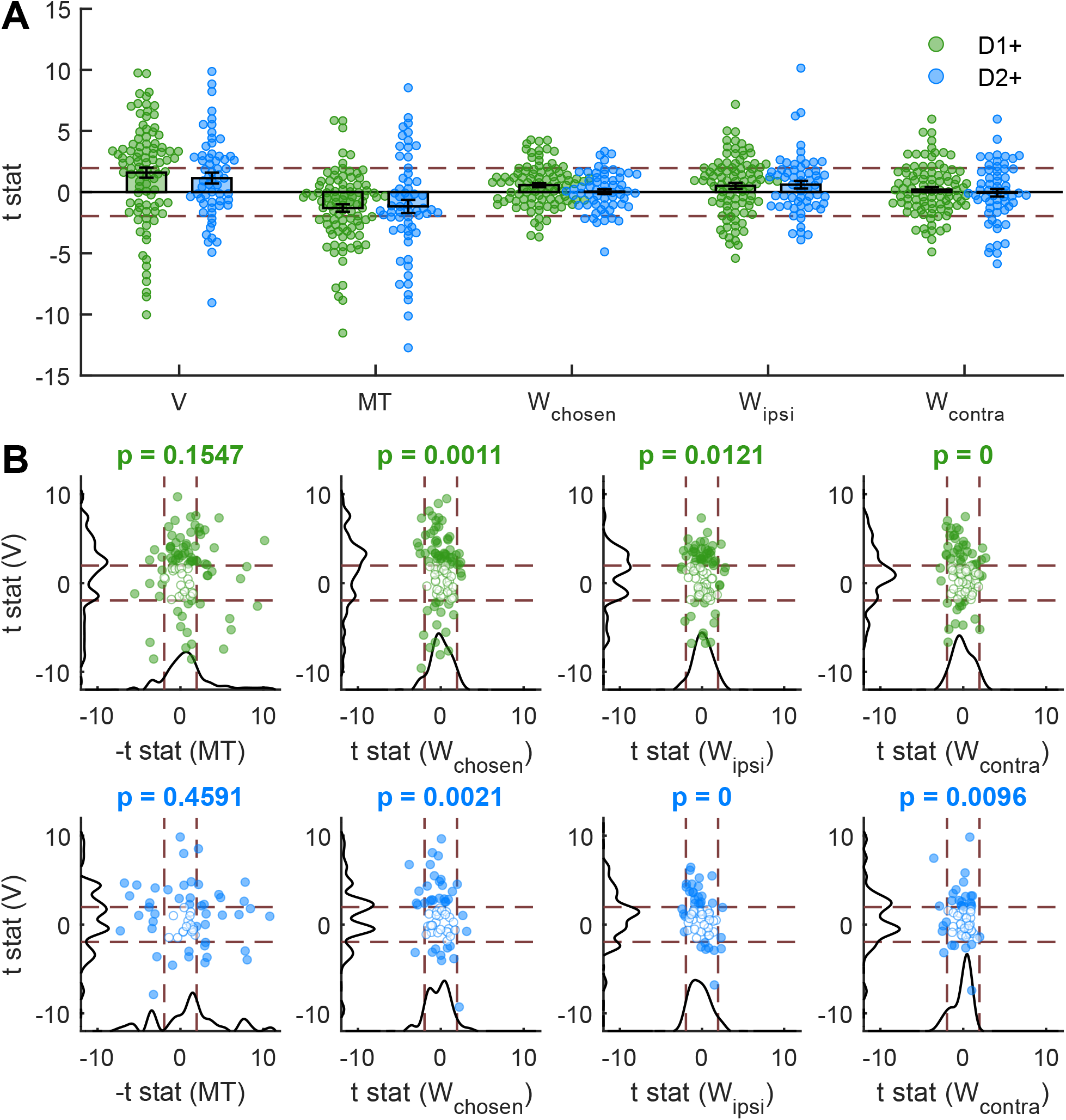
Both D1+ and D2+ SPNs signal overall trial value rather than specific action values during choice execution. **A**, Relationships between individual D1+ and D2+ cell firing during choice execution and a range of potential regressors: V – trial value, MT – Movement Time (Center-Out to Side-In duration, (log10)), W_*chosen*_ – chosen action weight, W_*ipsi*_ – ipsiversive action weight, W_*contra*_ – contraversive action weight. Here regressions for each factor were performed separately. **B**, t statistics for multiple regressions of D1+ (green) and D2+ (blue) responses during action selection (200 ms prior to Side-In). Dashed lines indicate significance thresholds (t = *±*.96). Distribution histograms for regressors on each axis are in black. p values for permutation tests between regressor t statistics are above panels in bold.

**Extended Data Figure 7.**
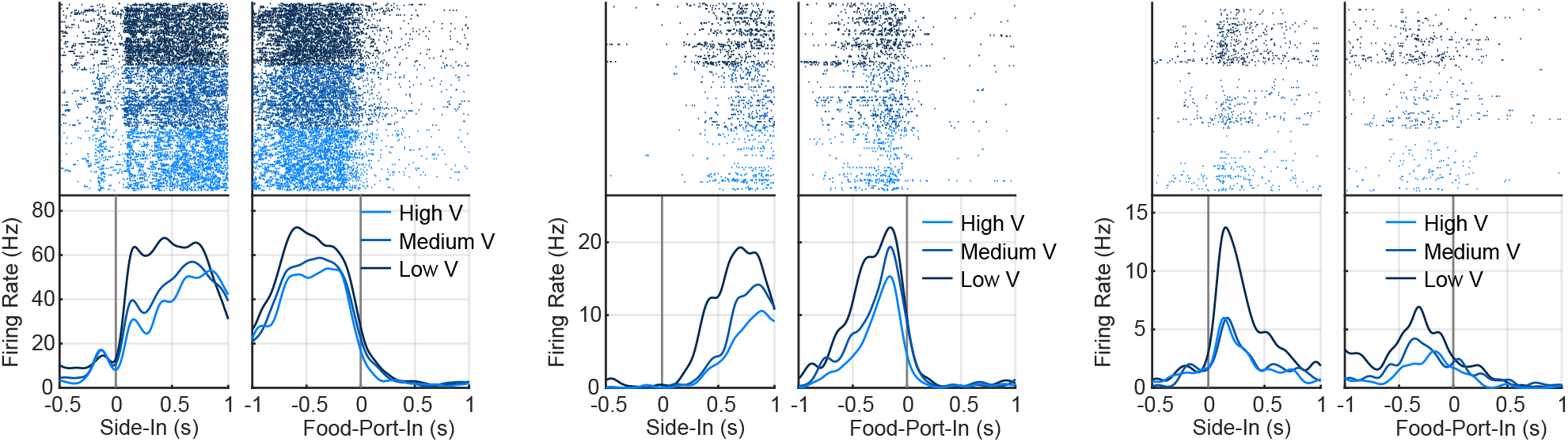
Examples of RPE-like responses in D2+ cells. Three example D2+ cells (left to right) each responding to rewarded outcomes with negative value scaling resembling RPE. Data are split into terciles of trial value (low, medium, high).

**Extended Data Figure 8.**
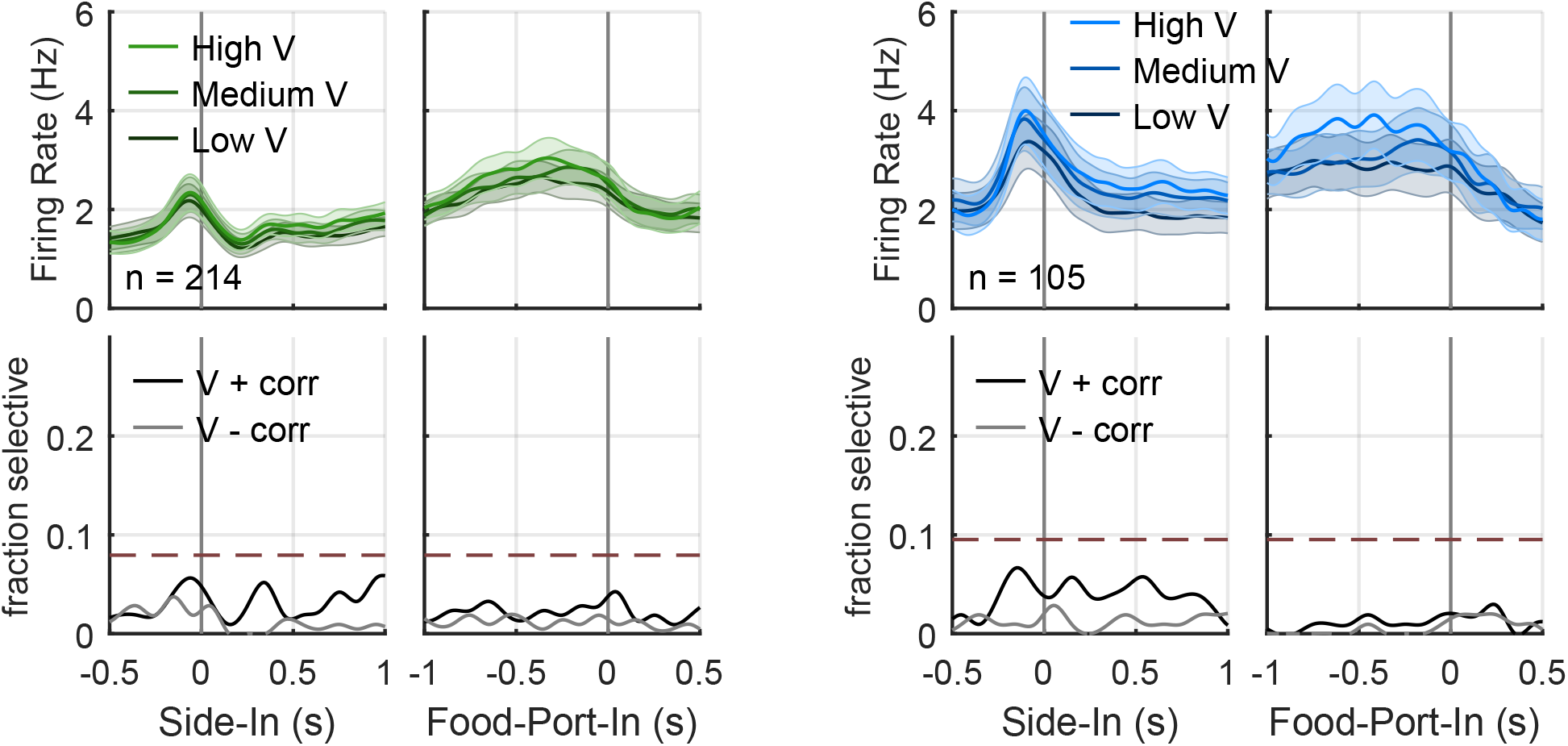
Lack of value signals in D1+ and D2+ cells following unrewarded outcomes. Mean firing of task-related D1+ and D2+ cells at Side-In and food port entry, split into terciles of trial value (V) for unrewarded trials (the main Fig. 6 shows data from rewarded trials). Bottom, fraction of D1+ SPNs at each moment whose firing has significant positive (V+) or negative (V-) correlations with value (p < 0.05, Pearson’s correlation, Bonferroni corrected). Dashed lines indicate when fractions are significantly higher than chance (binomial threshold, p < 0.05).

## Notes

### Competing Interest Statement

The authors have declared no competing interest.

## Bibliography

1. Peter W Kalivas and Nora D Volkow. The neural basis of addiction: a pathology of motivation and choice. American Journal of Psychiatry, 162(8):1403–1413, 2005.

2. Megan E Fox and Mary Kay Lobo. The molecular and cellular mechanisms of depression: a focus on reward circuitry. Molecular psychiatry, 24(12):1798–1815, 2019.

3. Tao R Wang, Sofie Moosa, Robert F Dallapiazza, W Jeffrey Elias, and William J Lynch. Deep brain stimulation for the treatment of drug addiction. Neurosurgical focus, 45(2):E11, 2018.

4. Katherine E Savell, Jennifer J Tuscher, Matthew E Zipperly, Charles G Duke, Robert A Phillips III, Ashley J Bauman, …, and Jeremy J Day. A dopamine-induced gene expression signature regulates neuronal function and cocaine response. Science advances, 6(26):eaba4221, 2020.

5. Jesus Bertran-Gonzalez, Clémentine Bosch, Matthieu Maroteaux, Miriam Matamales, Denis Hervé, Emmanuel Valjent, and Jean-Antoine Girault. Opposing patterns of signaling activation in dopamine d1 and d2 receptor-expressing striatal neurons in response to cocaine and haloperidol. Journal of Neuroscience, 28(22):5671–5685, 2008.

6. Satoshi Ikemoto and Jaak Panksepp. The role of nucleus accumbens dopamine in motivated behavior: a unifying interpretation with special reference to reward-seeking. Brain Research Reviews, 31(1):6–41, 1999.

7. John D Salamone and Merce Correa. The mysterious motivational functions of mesolimbic dopamine. Neuron, 76(3):470–485, 2012.

8. Joshua D Berke. What does dopamine mean? Nature neuroscience, 21(6):787–793, 2018.

9. J Todd Moyer, John A Wolf, and Leif H Finkel. Effects of dopaminergic modulation on the integrative properties of the ventral striatal medium spiny neuron. Journal of neurophysiology, 98(6):3731–3748, 2007.

10. Abhishek K Lahiri and Mark D Bevan. Dopaminergic transmission rapidly and persistently enhances excitability of d1 receptor-expressing striatal projection neurons. Neuron, 106 (2):277–290, 2020.

11. Salvador Hernandez-Lopez, Tatiana Tkatch, Enrique Perez-Garci, Elvira Galarraga, Jose Bargas, Heidi Hamm, and D James Surmeier. D2 dopamine receptors in striatal medium spiny neurons reduce l-type ca2+ currents and excitability via a novel plcβ1–ip3–calcineurin-signaling cascade. Journal of Neuroscience, 20(24):8987–8995, 2000.

12. Anastasia Klaus, Juliana Alves da Silva, and Rui M Costa. What, if, and when to move: basal ganglia circuits and self-paced action initiation. Annual review of neuroscience, 42: 459–483, 2019.

13. Alexxai V Kravitz, Bradley S Freeze, Philip R Parker, Kelsey Kay, Myo T Thwin, Karl Deisseroth, and Anatol C Kreitzer. Regulation of parkinsonian motor behaviours by optogenetic control of basal ganglia circuitry. Nature, 466(7306):622–626, 2010.

14. Wen Shen, Marc Flajolet, Paul Greengard, and D James Surmeier. Dichotomous dopaminergic control of striatal synaptic plasticity. Science, 321:848–851, 2008.

15. Wolfram Schultz, Peter Dayan, and P Read Montague. A neural substrate of prediction and reward. Science, 275(5306):1593–1599, 1997.

16. Neir Eshel, Jing Tian, Matthew Bukwich, and Naoshige Uchida. Dopamine neurons share common response function for reward prediction error. Nature neuroscience, 19(3):479–486, 2016.

17. Ali Mohebi, Jeffrey R Pettibone, Arif A Hamid, Jamie MT Wong, Lauren T Vinson, Tommaso Patriarchi, …, and Joshua D Berke. Dissociable dopamine dynamics for learning and motivation. Nature, 570(7759):65–70, 2019.

18. Richard S Sutton and Andrew G Barto. Reinforcement Learning: An Introduction. MIT press, 2018.

19. Joshua D Berke and Steven E Hyman. Addiction, dopamine, and the molecular mechanisms of memory. Neuron, 25(3):515–532, 2000.

20. Sho Yagishita, Akiko Hayashi-Takagi, Graham C Ellis-Davies, Hidetoshi Urakubo, Shunichi Ishii, and Haruo Kasai. A critical time window for dopamine actions on the structural plasticity of dendritic spines. Science, 345(6204):1616–1620, 2014.

21. Yuki Iino, Tsuyoshi Sawada, Kazuhiko Yamaguchi, Masaki Tajiri, Shin Ishii, Haruo Kasai, and Sho Yagishita. Dopamine d2 receptors in discrimination learning and spine enlargement. Nature, 579(7800):555–560, 2020.

22. Ameya Jaskir and Michael J Frank. On the normative advantages of dopamine and striatal opponency for learning and choice. Elife, 12:e85107, 2023.

23. Carina Soares-Cunha, Bárbara Coimbra, Ana David-Pereira, Sandra Borges, Luisa Pinto, Patrícia Costa, …, and Ana João Rodrigues. Activation of d2 dopamine receptor-expressing neurons in the nucleus accumbens increases motivation. Nature Communications, 7(1):11829, 2016.

24. Carina Soares-Cunha, Nuno A de Vasconcelos, Bárbara Coimbra, Ana V Domingues, João M Silva, Elsa Loureiro-Campos, …, and Ana João Rodrigues. Nucleus accumbens medium spiny neurons subtypes signal both reward and aversion. Molecular Psychiatry, 25(12):3241–3255, 2020.

25. Wolfram Schultz, Paul Apicella, Eugenio Scarnati, and Tomas Ljungberg. Neuronal activity in monkey ventral striatum related to the expectation of reward. Journal of Neuroscience, 12(12):4595–4610, 1992.

26. Saleem M Nicola, Irene A Yun, Ken T Wakabayashi, and Howard L Fields. Cue-evoked firing of nucleus accumbens neurons encodes motivational significance during a discriminative stimulus task. Journal of neurophysiology, 91(4):1840–1865, 2004.

27. Jeremy J Day, Jamie D Roitman, R Mark Wightman, and Regina M Carelli. Nucleus accumbens neurons differentially encode motivation and reinforcement. Journal of Neuroscience, 31(18):5876–5881, 2011.

28. Makoto Ito and Kenji Doya. Distinct neural representation in the dorsolateral, dorsomedial, and ventral parts of the striatum during fixed-and free-choice tasks. Journal of Neuroscience, 35(8):3499–3514, 2015.

29. Biao Tan, Theresa Nöbauer, Caitlin J Browne, Eric J Nestler, Alipasha Vaziri, and Jeffrey M Friedman. Dynamic processing of hunger and thirst by common mesolimbic neural ensembles. Proceedings of the National Academy of Sciences, 119(43):e2211688119, 2022.

30. Tomomi Nishioka, Supawat Attachaipanich, Kenji Hamaguchi, Michael Lazarus, Alban de Kerchove d’Exaerde, Tom Macpherson, and Takatoshi Hikida. Error-related signaling in nucleus accumbens d2 receptor-expressing neurons guides inhibition-based choice behavior in mice. Nature Communications, 14(1):2284, 2023.

31. Shinya Nonomura, Kazuki Nishizawa, Yutaka Sakai, Yasuo Kawaguchi, Sho Kato, Motokazu Uchigashima, …, and Minoru Kimura. Monitoring and updating of action selection for goal-directed behavior through the striatal direct and indirect pathways. Neuron, 99(6): 1302–1314, 2018.

32. Jia H Shin, Daejong Kim, and Min W Jung. Differential coding of reward and movement information in the dorsomedial striatal direct and indirect pathways. Nature Communications, 9(1):404, 2018.

33. Arif A Hamid, Jeffrey R Pettibone, Omar S Mabrouk, Victoria L Hetrick, Rebecca Schmidt, Caitlin M Vander Weele, and Joshua D Berke. Mesolimbic dopamine signals the value of work. Nature neuroscience, 19(1):117–126, 2016.

34. Ilhong A Yun, Saleem M Nicola, and Howard L Fields. Contrasting effects of dopamine and glutamate receptor antagonist injection in the nucleus accumbens suggest a neural mechanism underlying cue-evoked goal-directed behavior. European Journal of Neuroscience, 20(1):249–263, 2004.

35. Jeffrey R Pettibone, Yinying Y Jai, Regina C Derman, Timothy W Faust, Elizabeth D Hughes, William E Filipiak, …, and Joshua D Berke. Knock-in rat lines with cre recombinase at the dopamine d1 and adenosine 2a receptor loci. eneuro, 6:5, 2019.

36. Adel A Alcantara, Victor Chen, Bruce E Herring, Jill M Mendenhall, and Maria L Berlanga. Localization of dopamine d2 receptors on cholinergic interneurons of the dorsal striatum and nucleus accumbens of the rat. Brain research, 986(1-2):22–29, 2003.

37. Pierre F Durieux, Bertrand Bearzatto, Stefania Guiducci, Thorsten Buch, Ari Waisman, Michele Zoli, Serge N Schiffmann, and Alban de Kerchove d’Exaerde. D2r striatopallidal neurons inhibit both locomotor and drug reward processes. Nature neuroscience, 12(4): 393–395, 2009.

38. Nathan C Klapoetke, Yukiko Murata, Soo Yeun Kim, Stefan R Pulver, Anna Birdsey-Benson, Yong Ku Cho, and Edward S Boyden. Independent optical excitation of distinct neural populations. Nature methods, 11(3):338–346, 2014.

39. Sara Ghods-Sharifi and Stan B Floresco. Differential effects on effort discounting induced by inactivations of the nucleus accumbens core or shell. Behavioral neuroscience, 124(2): 179, 2010.

40. Joshua D Berke, Murat Okatan, Jennifer Skurski, and Howard B Eichenbaum. Oscillatory entrainment of striatal neurons in freely moving rats. Neuron, 43(6):883–896, 2004.

41. C S Lansink, P M Goltstein, J V Lankelma, and C M Pennartz. Fast-spiking interneurons of the rat ventral striatum: Temporal coordination of activity with principal cells and responsiveness to reward. European Journal of Neuroscience, 32(3):494–508, 2010.

42. Gregory J Gage, Colin R Stoetzner, Alexander B Wiltschko, and Joshua D Berke. Selective activation of striatal fast-spiking interneurons during choice execution. Neuron, 67(3):466–479, 2010.

43. Sharif A Taha and Howard L Fields. Inhibitions of nucleus accumbens neurons encode a gating signal for reward-directed behavior. Journal of Neuroscience, 26(1):217–222, 2006.

44. Kazuyuki Samejima, Yoshiyuki Ueda, Kenji Doya, and Minoru Kimura. Representation of action-specific reward values in the striatum. Science, 310(5752):1337–1340, 2005.

45. Hyojung Kim, Ji Hoon Sul, Namjung Huh, Daeyeol Lee, and Min W Jung. Role of striatum in updating values of chosen actions. Journal of neuroscience, 29(47):14701–14712, 2009.

46. Eun Jung Shin, Yunji Jang, Soyeon Kim, Hyojung Kim, Xiaodong Cai, Hyojung Lee, …, and Min W Jung. Robust and distributed neural representation of action values. Elife, 10: e53045, 2021.

47. Michael J Frank. Dynamic dopamine modulation in the basal ganglia: a neurocomputational account of cognitive deficits in medicated and nonmedicated parkinsonism. Journal of cognitive neuroscience, 17(1):51–72, 2005.

48. Kei Oyama, István Hernádi, Toshio Iijima, and Ken-Ichiro Tsutsui. Reward prediction error coding in dorsal striatal neurons. Journal of neuroscience, 30(34):11447–11457, 2010.

49. GJ Mogenson, DL Jones, and CY Yim. From motivation to action: functional interface between the limbic system and the motor system. Progress in neurobiology, 14(2-3):69–97, 1980.

50. Trevor W Robbins and Barry J Everitt. Neurobehavioural mechanisms of reward and motivation. Current opinion in neurobiology, 6(2):228–236, 1996.

51. Brian A Baldo and Ann E Kelley. Discrete neurochemical coding of distinguishable motivational processes: insights from nucleus accumbens control of feeding. Psychopharmacology, 191:439–459, 2007.

52. Mathilde Sicre, Julie Meffre, Didier Louber, and Frédéric Ambroggi. The nucleus accumbens core is necessary for responding to incentive but not instructive stimuli. Journal of Neuroscience, 40(6):1332–1343, 2020.

53. June Y Chang, Susan F Sawyer, Robert S Lee, and Donald J Woodward. Electrophysiological and pharmacological evidence for the role of the nucleus accumbens in cocaine self-administration in freely moving rats. Journal of neuroscience, 14(3):1224–1244, 1994.

54. Jeremy J Day, Robert A Wheeler, Mitchell F Roitman, and Regina M Carelli. Nucleus accumbens neurons encode pavlovian approach behaviors: evidence from an autoshaping paradigm. European Journal of Neuroscience, 23(5):1341–1351, 2006.

55. Franscesco Ambroggi, Akinori Ishikawa, Howard L Fields, and Saleem M Nicola. Basolateral amygdala neurons facilitate reward-seeking behavior by exciting nucleus accumbens neurons. Neuron, 59(4):648–661, 2008.

56. Matthijs AA van der Meer, Adam Johnson, Neil C Schmitzer-Torbert, and A David Redish. Triple dissociation of information processing in dorsal striatum, ventral striatum, and hippocampus on a learned spatial decision task. Neuron, 67(1):25–32, 2010.

57. Erin S Calipari, Rosemary C Bagot, Immanuel Purushothaman, Thomas J Davidson, Jordan T Yorgason, Catherine J Peña, …, and Eric J Nestler. In vivo imaging identifies temporal signature of d1 and d2 medium spiny neurons in cocaine reward. Proceedings of the National Academy of Sciences, 113(10):2726–2731, 2016.

58. Timothy J O’Neal, Michael X Bernstein, Douglas J MacDougall, and Susan M Ferguson. A conditioned place preference for heroin is signaled by increased dopamine and direct pathway activity and decreased indirect pathway activity in the nucleus accumbens. Journal of Neuroscience, 42(10):2011–2024, 2022.

59. Jacqueline Du Hoffmann and Saleem M Nicola. Dopamine invigorates reward seeking by promoting cue-evoked excitation in the nucleus accumbens. Journal of Neuroscience, 34 (43):14349–14364, 2014.

60. Kent C Berridge. The debate over dopamine’s role in reward: the case for incentive salience. Psychopharmacology, 191:391–431, 2007.

61. Saleem M Nicola. The flexible approach hypothesis: unification of effort and cue-responding hypotheses for the role of nucleus accumbens dopamine in the activation of reward-seeking behavior. Journal of Neuroscience, 30(49):16585–16600, 2010.

62. Ali Mohebi, Val L Collins, and Joshua D Berke. Accumbens cholinergic interneurons dynamically promote dopamine release and enable motivation. eLife, 12:e85011, 2023.

63. Brenda B Gore and Larry S Zweifel. Genetic reconstruction of dopamine d1 receptor signaling in the nucleus accumbens facilitates natural and drug reward responses. Journal of Neuroscience, 33(20):8640–8649, 2013.

64. Rosario Moratalla, Mei Xu, Susumu Tonegawa, and Ann M Graybiel. Cellular responses to psychomotor stimulant and neuroleptic drugs are abnormal in mice lacking the d1 dopamine receptor. Proceedings of the National Academy of Sciences, 93(25):14928–14933, 1996.

65. Joshua D Berke, Ronald F Paletzki, Geoffrey J Aronson, Steven E Hyman, and Charles R Gerfen. A complex program of striatal gene expression induced by dopaminergic stimulation. Journal of Neuroscience, 18(14):5301–5310, 1998.

66. Yuki Nakamura, Sophie Longueville, Akinori Nishi, Denis Hervé, Jean-Antoine Girault, and Yusuke Nakamura. Dopamine d1 receptor-expressing neurons activity is essential for locomotor and sensitizing effects of a single injection of cocaine. European Journal of Neuroscience, 54(4):5327–5340, 2021.

67. Wolfram Schultz. Multiple dopamine functions at different time courses. Annual Review of Neuroscience, 30:259–288, 2007.

68. Robert Lindroos, Matthijs C Dorst, Kai Du, Marko Filipović, Daniel Keller, Mira Ketzef, …, and Jeanette Hellgren Kotaleski. Basal ganglia neuromodulation over multiple temporal and structural scales—simulations of direct pathway msns investigate the fast onset of dopaminergic effects and predict the role of kv4.2. Frontiers in neural circuits, 12:3, 2018.

69. João António da Silva, Francisco Tecuapetla, Vîtor Paixão, and Rui M Costa. Dopamine neuron activity before action initiation gates and invigorates future movements. Nature, 554:244–248, 2018.

70. Mitchell F Roitman, Garret D Stuber, Paul E Phillips, R Mark Wightman, and Regina M Carelli. Dopamine operates as a subsecond modulator of food seeking. Journal of Neuroscience, 24(6):1265–1271, 2004.

71. Matthew W Howe, Patrick L Tierney, Sharon G Sandberg, Paul E Phillips, and Ann M Graybiel. Prolonged dopamine signalling in striatum signals proximity and value of distant rewards. Nature, 500(7464):575–579, 2013.

72. Talia Krausz, Allison E Comrie, Andrea E Kahn, Loren M Frank, Nathaniel D Daw, and Joshua D Berke. Dual credit assignment processes underlie dopamine signals in a complex spatial environment. Neuron, 2023. In press.

73. Long Ding and David J Perkel. Dopamine modulates excitability of spiny neurons in the avian basal ganglia. Journal of Neuroscience, 22(12):5210–5218, 2002.

74. Jesper Ericsson, Marcus Stephenson-Jones, Juan Pérez-Fernández, Birgit Robertson, Gilad Silberberg, and Sten Grillner. Dopamine differentially modulates the excitability of striatal neurons of the direct and indirect pathways in lamprey. Journal of Neuroscience, 33(18):8045–8054, 2013.

75. Eric K Richfield, John B Penney, and Anne B Young. Anatomical and affinity state comparisons between dopamine d1 and d2 receptors in the rat central nervous system. Neuroscience, 30(3):767–777, 1989.

76. Pamela F Marcott, Aphroditi A Mamaligas, and Christopher P Ford. Phasic dopamine release drives rapid activation of striatal d2-receptors. Neuron, 84(1):164–176, 2014.

77. Jakob K Dreyer, Kjartan F Herrik, Rune W Berg, and Jørn D Hounsgaard. Influence of phasic and tonic dopamine release on receptor activation. Journal of Neuroscience, 30 (42):14273–14283, 2010.

78. Ching Liu, Pragya S Goel, and Pascal S Kaeser. Spatial and temporal scales of dopamine transmission. Nature Reviews Neuroscience, 22(6):345–358, 2021.

79. Sarah L Cole, Mike J Robinson, and Kent C Berridge. Optogenetic self-stimulation in the nucleus accumbens: D1 reward versus d2 ambivalence. PLoS One, 13(11):e0207694, 2018.

80. Moritz Möller and Rafal Bogacz. Learning the payoffs and costs of actions. PLoS computational biology, 15(2):e1006285, 2019.

81. John NJ Reynolds, Brian I Hyland, and Jeff R Wickens. A cellular mechanism of reward-related learning. Nature, 413(6851):67–70, 2001.

82. Kevin N Gurney, Mark D Humphries, and Peter Redgrave. A new framework for cortico-striatal plasticity: behavioural theory meets in vitro data at the reinforcement-action interface. PLoS biology, 13(1):e1002034, 2015.

83. Wulfram Gerstner, Mike Lehmann, Vasiliki Liakoni, Derek Corneil, and Johanni Brea. Eligibility traces and plasticity on behavioral time scales: experimental support of neohebbian three-factor learning rules. Frontiers in neural circuits, 12:53, 2018.

84. Guohong Cui, Sang Beom Jun, Xin Jin, Mimi D Pham, Steven S Vogel, David M Lovinger, and Rui M Costa. Concurrent activation of striatal direct and indirect pathways during action initiation. Nature, 494(7436):238–242, 2013.

85. Jeffrey E Markowitz, William F Gillis, Chris C Beron, Sharon Q Neufeld, Keiramarie Robertson, Nikhil D Bhagat, …, and Subimal Ray Datta. The striatum organizes 3d behavior via moment-to-moment action selection. Cell, 174(1):44–58, 2018.

86. Justus G Parker, John D Marshall, Buz Ahanonu, Yi-Wen Wu, Tae H Kim, Benjamin F Grewe, …, and Mark J Schnitzer. Diametric neural ensemble dynamics in parkinsonian and dyskinetic states. Nature, 557(7704):177–182, 2018.

87. Fatuel Tecuapetla, Stephanie Matias, Guillaume P Dugue, Zachary F Mainen, and Ricardo M Costa. Balanced activity in basal ganglia projection pathways is critical for contraversive movements. Nature Communications, 5(1):4315, 2014.

88. Makoto Ito and Kenji Doya. Validation of decision-making models and analysis of decision variables in the rat basal ganglia. Journal of Neuroscience, 29(31):9861–9874, 2009.

89. Okihide Hikosaka, Kenji Nakamura, and Hiromichi Nakahara. Basal ganglia orient eyes to reward. Journal of neurophysiology, 95(2):567–584, 2006.

90. Nathan F Parker, Courtney M Cameron, Joshua P Taliaferro, Jeongyeon Lee, Jaewon Y Choi, Thomas J Davidson, …, and Ilana B Witten. Reward and choice encoding in terminals of midbrain dopamine neurons depends on striatal target. Nature neuroscience, 19(6):845–854, 2016.

91. M Mallar Moss, Peter Zatka-Haas, Kenneth D Harris, Matteo Carandini, and Armin Lak. Dopamine axons in dorsal striatum encode contralateral visual stimuli and choices. Journal of Neuroscience, 41(34):7197–7205, 2021.

92. Reiko Kawagoe, Yoriko Takikawa, and Okihide Hikosaka. Expectation of reward modulates cognitive signals in the basal ganglia. Nature neuroscience, 1(5):411–416, 1998.

93. Xinying Cai, Soo Hyun Kim, and Daeyeol Lee. Heterogeneous coding of temporally discounted values in the dorsal and ventral striatum during intertemporal choice. Neuron, 69(1):170–182, 2011.

94. Brian L Goldstein, Brian R Barnett, Gabriel Vasquez, Steven C Tobia, Vadim Kashtelyan, Amanda C Burton, and Matthew R Roesch. Ventral striatum encodes past and predicted value independent of motor contingencies. Journal of Neuroscience, 32(6):2027–2036, 2012.

95. Bruno Averbeck and John P O’Doherty. Reinforcement-learning in fronto-striatal circuits. Neuropsychopharmacology, 47(1):147–162, 2022.

96. Seong-Whan Hong and Okihide Hikosaka. Dopamine-mediated learning and switching in cortico-striatal circuit explain behavioral changes in reinforcement learning. Frontiers in behavioral neuroscience, 5:15, 2011.

97. Yoshikazu Isomura, Takashi Takekawa, Rie Harukuni, Takashi Handa, Hidenori Aizawa, Masahiko Takada, and Tomoki Fukai. Reward-modulated motor information in identified striatum neurons. Journal of Neuroscience, 33(25):10209–10220, 2013.

98. Oscar Bartra, Joseph T McGuire, and Joseph W Kable. The valuation system: a coordinate-based meta-analysis of bold fmri experiments examining neural correlates of subjective value. Neuroimage, 76:412–427, 2013.

99. David M Smith and M M Torregrossa. Valence encoding in the amygdala influences motivated behavior. Behavioural Brain Research, 411:113370, 2021.

100. Jane X Wang, Zeb Kurth-Nelson, Dharshan Kumaran, Dileep Tirumala, Hubert Soyer, Joel Z Leibo, …, and Matthew Botvinick. Prefrontal cortex as a meta-reinforcement learning system. Nature Neuroscience, 21(6):860–868, 2018.

101. R. Corey Evans, Evan L. Twedell, Michael Zhu, Jack Ascencio, Ruibin Zhang, and Zayd M. Khaliq. Functional dissection of basal ganglia inhibitory inputs onto substantia nigra dopaminergic neurons. Cell reports, 32(11), 2020.

102. Mark A Farries, Timothy W Faust, Ali Mohebi, and Joshua D Berke. Selective encoding of reward predictions and prediction errors by globus pallidus subpopulations. Current Biology, 2023. In press.

103. Jonathan W Mink. The basal ganglia: focused selection and inhibition of competing motor programs. Progress in neurobiology, 50(4):381–425, 1996.

104. Fatuel Tecuapetla, Xin Jin, Suzana Q Lima, and Ricardo M Costa. Complementary contributions of striatal projection pathways to action initiation and execution. Cell, 166 (3):703–715, 2016.

105. Jeffrey W Dalley, Tim D Fryer, Laurent Brichard, Emma S Robinson, David E Theobald, Kaisa Lääne, …, and Trevor W Robbins. Nucleus accumbens d2/3 receptors predict trait impulsivity and cocaine reinforcement. Science, 315(5816):1267–1270, 2007.

106. Jason E Chung, Jeremy F Magland, Alex H Barnett, Vanessa M Tolosa, Angela C Tooker, Kye Y Lee, Kedar G Shah, Sarah H Felix, Loren M Frank, and Leslie F Greengard. A fully automated approach to spike sorting. Neuron, 95(6):1381–1394, 2017.

107. Liora Elber-Dorozko and Yonatan Loewenstein. Striatal action-value neurons reconsidered. Elife, 7:e34248, 2018.

